# A family of metabolite damage-control phosphatases modulates ROS-induced autophagy and plant stress resilience

**DOI:** 10.64898/2026.09.29.755255

**Authors:** Florentine Ballhaus, Viviana Vella, Yong Zou, Glenn R Hicks, Jacob Whittaker, Adrian Suarez Covarrubias, Flavio Ballante, Ali Moazzami, Igor Sabljic, Mary A Nishiotis, Anette Roos, Selvaraju Kanagarajan, Stina Berglund-Fick, Weixing Qian, Gustav Nestor, Peter V Bozhkov, Elena A Minina, Adrian N Dauphinee

## Abstract

Autophagy is an intracellular recycling pathway with profound impacts on development, growth, and stress tolerance in eukaryotes. Therefore, unravelling signalling mechanisms that modulate the process has broad applicability. Here, we characterised a small organic molecule as an enhancer of autophagy across diverse plant lineages. Through an *in vivo* photoaffinity proteomics approach in *Arabidopsis thaliana* seedlings, we identified Domain of Unknown Function 89 (DUF89) proteins as molecular targets. Hereafter, we refer to these proteins as Reactive Metabolite Damage-Control Phosphatases (RMDPs), which are metabolite-repair phosphatases that clear reactive products. The new autophagy modulator is called RMDP inhbitor-1 (RMDPi-1) due to its demonstrated inhibitory effect on AtRMDP1 activity *in vitro.* Binding was further confirmed through co-incubation that thermostabilised the protein. Crystal structures of AtRMDP1 were obtained via X-ray diffraction, confirming binding of RMDPi-1 in the catalytic site and revealing a flexible region near the binding pocket that was absent in previous models. Molecular docking predicted two possible binding modes consistent with the observed electron density. RMDPi-1 treatment and the phosphatase gene knockout increased autophagic flux in Arabidopsis and *Chlamydomonas reinhardtii,* establishing a conserved effect. The observed autophagy response is linked to a spike in reactive oxygen species (ROS). *AtRMDP* knockout also results in higher ROS tolerance and greater biomass in Arabidopsis seedlings. Targeted and untargeted metabolomic experiments revealed that both pharmacological and genetic suppression of RMDP leads to accumulation of glycating agents and phosphate sugars, associating loss of RMDP activity to the ROS stress that induces alternate ROS-scavenging mechanisms, and autophagy. Taken together, our integrated chemical genetics approach reveals enhanced autophagy via inhibition of RMDP family members, which impacts on plant stress tolerance via conserved metabolite damage-control functions.

## Introduction

Autophagy is a conserved catabolic process that maintains cellular homeostasis by sequestering damaged or obsolete components within a double-membrane autophagosome for delivery to the lysosome (in animals) or lytic vacuole (in fungi and plants) for subsequent degradation and recycling (Marshall & Vierstra 2018; Su et al. 2020). Autophagy operates at two levels: under basal conditions, it acts as a housekeeping quality-control mechanism, whereas during stress and starvation autophagy is strongly upregulated, driving extensive recycling of cellular components and reducing the cell’s dependence on external nutrient input (Rabinowitz & White 2010).

Given its fundamental role in cellular health and stress adaptation, autophagy has attracted considerable interest in plant and animal systems, although research in plants has lagged despite its direct impact on plant fitness. *Arabidopsis thaliana* plants overexpressing autophagy-related *(ATG)* genes exhibit heightened stress tolerance (Liu et al. 2009), increased fecundity, and improved yield (Avin-Wittenberg et al. 2018; Minina et al. 2018), whereas knockouts of the same genes have the opposite effect (Minina et al. 2018). Constant stimulation of autophagy may adversely affect plant fitness, as upregulated autophagic flux can be exploited by plant viruses and virulent bacteria, making plants more susceptible to diseases (Chen et al. 2017; Ustun et al. 2018). In terms of increased plant fitness, this observation emphasises the value of transiently upregulating autophagy. For example, short-term treatments that enhance autophagic flux could benefit agricultural plants facing abiotic stresses like drought. Such transient control requires chemical autophagy effectors that can be easily modulated with low cytotoxicity, which we currently lack. For example, the TOR (target of rapamycin) kinase inhibitor, AZD8055, strongly enhances autophagic flux in plants and mammalian cells (Chresta et al. 2010; Dong et al. 2015); however, treating plants with AZD8055 leads to stunted growth. Other available modulators are similarly cytotoxic, as exemplified by the PI3K inhibitor 3-ethyladenine (3-MA), which both suppresses and, on prolonged treatment, unexpectedly enhances autophagy (Wu et al. 2010; Klionsky et al. 2021).

To identify modulable plant autophagy effectors, we developed a multiphasic chemical screening pipeline that revealed several promising small organic molecules that are effective in tobacco BY-2 cell cultures and Arabidopsis seedlings (Dauphinee et al. 2019). Importantly, these compounds do not adversely affect other endomembrane system components, distinguishing them from available autophagy modulators and warranting their further development. Here, we characterised one of our autophagy enhancers through an integrative chemical genetics and multi-omics approach (Cong et al. 2012). Using *in vivo* photoaffinity labelling and proteomics, we identified domain of unknown function 89 (DUF89) phosphatases as molecular targets. The DUF89 family includes a conserved clade of metal-dependent phosphatases and is represented across taxa and all domains of life. There are three DUF89 subfamilies. The one relevant here, class II, has representatives in animals, fungi, and plants that occur either as pantothenate kinase (PANK)-DUF89 fusion proteins or stand-alone phosphatases. These enzymes are thought to function in metabolite damage control by dephosphorylating potentially cytotoxic phosphometabolites (Huang et al. 2016; Griffith et al. 2021). Hereafter, we refer to stand-alone DUF89 proteins as Reactive Metabolite Damage Phosphatases (RMDPs). Arabidopsis encodes only subfamily II proteins, including one PANK fusion variant, AtPANK2 (Tilton et al. 2006), and three stand-alone paralogs: AtRMDP1 (At2g17340), AtRMDP2 (At2g17320), and AtRMDP3 (At4g35360). The number of RMDPs varies among plant species. For example, the unicellular green alga *Chlamydomonas reinhardtii* encodes a single CrRDMP and a single CrPANK.

Research and interest in human RMDPs have recently increased due to the realisation that the human pseudokinase HsPANK4 (Yao et al. 2019) retains a critical, rate-limiting role in CoA biosynthesis via its functional DUF89 phosphatase domain (Dibble et al. 2022). Furthermore, HsPANK4 has physiological relevance in diseases including cancer (Vella et al. 2024), metabolic disorders (Miranda-Cervantes et al. 2025), and neurodegenerative disorders (Munshi et al. 2022; Hwang et al. 2025). In plants, RMDPs and their PANK-fusion counterparts are poorly understood.

We built on our initial discovery of novel plant autophagy modulators by discovering that AtRMDPs are the target of our autophagy enhancer, which we named RMPD inhibitor 1 (RMDPi-1). We demonstrate that RMDPi-1 directly binds to AtRMDP1 phosphatase and inhibits its activity *in vitro.* Both pharmacological inhibition by RMDPi-1 and genetic knockout in Arabidopsis and Chlamydomonas increase autophagic flux. Examining the physiological consequences, we find that RMDP loss or inhibition *in planta* perturbs central carbon metabolism and elevates reactive oxygen species (ROS), inducing autophagy. Through targeted and untargeted metabolomics of RMDPi-1-treated and RMDP-deficient Arabidopsis and Chlamydomonas, we detect the accumulation of phosphate sugars and potent glycating agents, along with ROS-scavenging metabolites. Together, our findings show that RMDPi-1 modulator of autophagic flux across evolutionarily distinct plant lineages and establish RMDP/DUF89 phosphatases as conserved metabolite damage-control proteins that regulate ROS-mediated autophagy.

## Results

### RMPDi-1 is an autophagy enhancer targeting DUF89 proteins

To identify protein targets of the novel autophagy enhancer RMDPi-1 (Figure 1a) in Arabidopsis, we developed a photoaffinity labelling (PAL) based pulldown assay (Smith & Collins 2015) (Figure 1c). The RMDPi-1 molecule was functionalised with a diazirine-based photoactivatable probe and designed based on structure-activity relationships (SARs). To define the SARs, we screened a library of RMDPi-1 analogs (Supplementary Files S1 and S2) for their effect on autophagic flux using the AuTToFlux tandem-tag (TT) assay (Dauphinee et al. 2020). In brief, the TT assay is a semi-automated confocal microscopy workflow that detects autophagic cargo delivery to the vacuole using an RFP-GFP-ATG8 tag. Since GFP fluorescence is quenched in acidic environments, increased vacuolar RFP fluorescence relative to GFP is indicative of increased autophagy levels. The SAR experiments revealed that 6-Oxo-1,6-dihydro-pyrimidine-5-carboxylic acid ethyl ester was most critical for RMDPi-1 functional activity. Thus, we designed and synthesised two PAL probes (Figure 1a) - an active RMDPi-1 probe that was expected to retain autophagy-enhancing activity, and a PAL-compatible control probe that should not induce autophagy.

**Figure 1.**
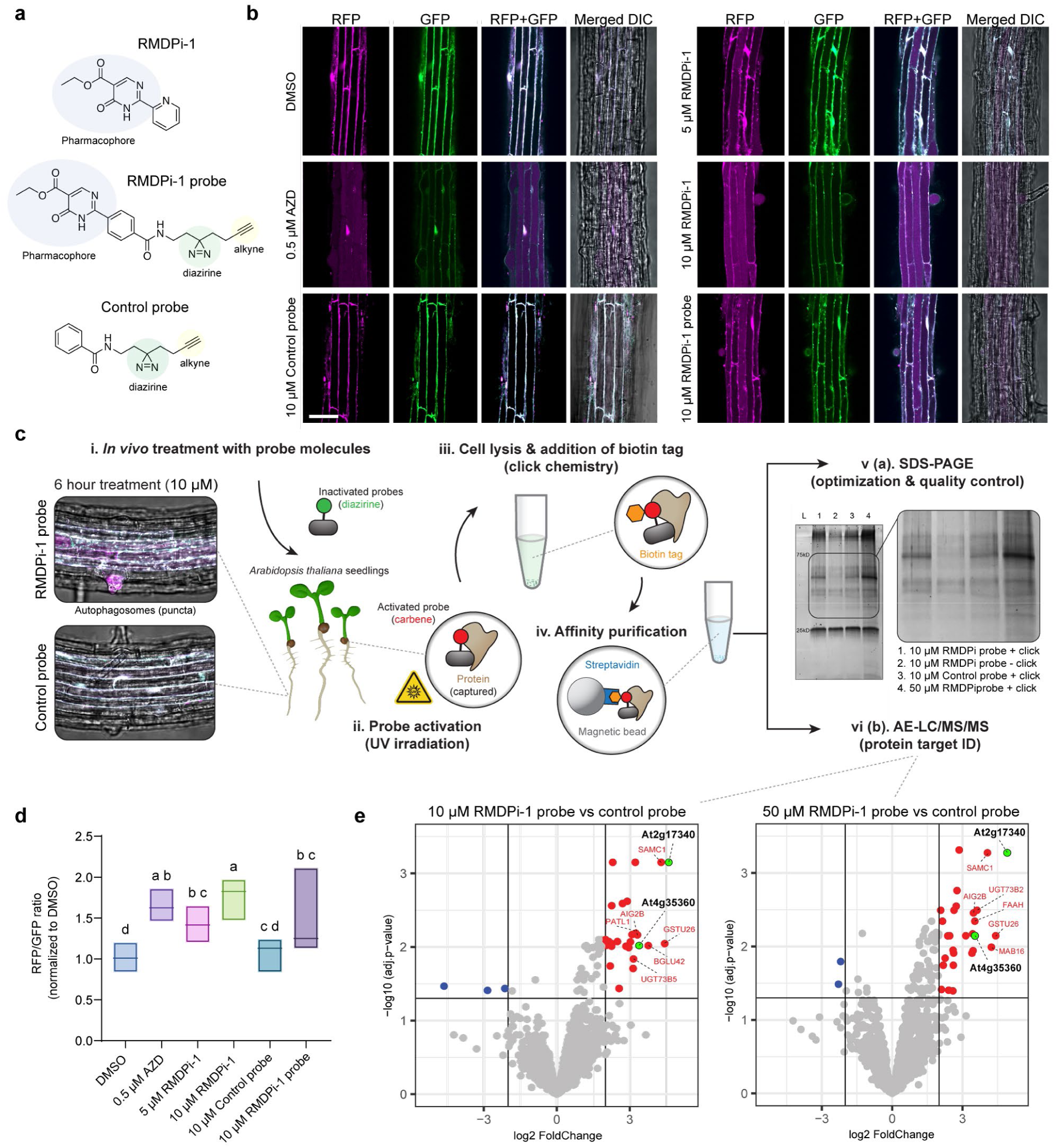
RMDPi-1 is an autophagy enhancer targeting DUF89 proteins. **a** Chemical structure of Reactive Metabolite Damage Phosphatase inhibitor 1 (RMDPi-1) highlighting the active pharmacophore (based on SARs), together with RMDPi-1 and control probes bearing diazirine and alkyne functional groups for the pulldown assay. **b** Confocal micrographs of 5-day-old Arabidopsis roots expressing the ATG8 TandemTag reporter (35S::TagRFP-mWasabi-ATG8a) in the Col-0 background. Seedlings were treated with DMSO (vehicle control), AZD8055 (positive control; autophagy inducer), RMDPi-1, RMDPi-1 probe, or negative control-probe. Scale bar: 20 µm. **c** Photoaffinity labelling (PAL) workflow for the identification of RMDPi-1 protein targets. Arabidopsis seedlings were treated for 6 h with either RMDPi-1 probe or the control-probe, followed by UV-induced probe activation. Cells were lysed, and probe-labelled proteins were conjugated to biotin via click chemistry. Proteins were enriched by affinity purification, and differential capture was validated by SDS-PAGE. Protein identities were determined by AE-LC-MS/MS. **d** Quantification of vacuolar RFP/GFP fluorescence ratios in Arabidopsis seedlings used for PAL assays. AZD8055, RMDPi-1 and RMDPi-1 probe treatment significantly increased autophagic flux compared to DMSO. The control-probe did not increase autophagy. One-way ANOVA, Tukey test, letters represent significantly different means, n ≥ 6 biological replicates. **e** Volcano plots of differential protein capture of RMDPi-1 probe vs. control-probe. Green circles mark enriched DUF89 protein family members. Red circles indicate all significantly enriched proteins, and blue circles are decreased proteins in the sample vs control. (n = 3 biological replicates; full proteomics data in Supplementary file S3).

TT assays demonstrated that treatment with RMDPi-1, as well as the RMDPi-1 probe and AZD8055 (a positive control for autophagy induction), significantly enhanced autophagic activity compared to the DMSO control (Figure 1b, d). In contrast, the structurally similar control probe had no detectable effect, confirming the suitability of the RMDPi-1 probe for downstream pulldown experiments. To identify putative protein targets, we performed affinity enrichment by LC-MS/MS and identified proteins that preferentially co-purified with the RMDPi-1 probe relative to the control probe (Figure 1e, Supplementary file S3). Among the significantly enriched proteins were two AtRMDP/DUF89 family members (Supplementary file S4): At2g17340 (AtRMDP1) and At4g35360 (AtRMDP3). Enrichment of both proteins increased with higher concentrations of the RMDPi-1 probe, with AtRMDP1 representing the top enriched candidate at the two probe concentrations used (Figure 1e). Together, the *in vivo* PAL workflow identified proteins from the DUF89 family as candidate targets of RMDPi-1.

### RMDPi-1 inhibits AtRMDP1 phosphatase activity via a structurally defined binding site

Because DUF89 proteins are metal-dependent phosphatases, we tested AtRMDP1 recombinant protein stability *in vitro* in the presence of metal cofactors and observed the highest stability with either Co^2^+ or Ni^2^+, consistent with previous biochemical characterisations (Huang et al. 2016). Binding of RMDPi-1 to AtRMDP1 was confirmed via thermal shift assay (TSA) and phosphatase activity assays (Figure 2a, b). Incubating AtRMDP1 with RMDPi-1 resulted in a shift of 5.7°C in the protein’s melting temperature *(T_m_)* relative to the DMSO control, indicating a strong thermal-stabilising effect. This stabilisation was dose-dependent, with a half-maximal effective concentration (EC50) of 97 µM (Figure 2a). Consistent with direct binding, RMDPi-1 reduced AtRMDP1 phosphatase activity in a dose-dependent manner, yielding an IC50 of 1.70 pM (Figure 2b).

**Figure 2.**
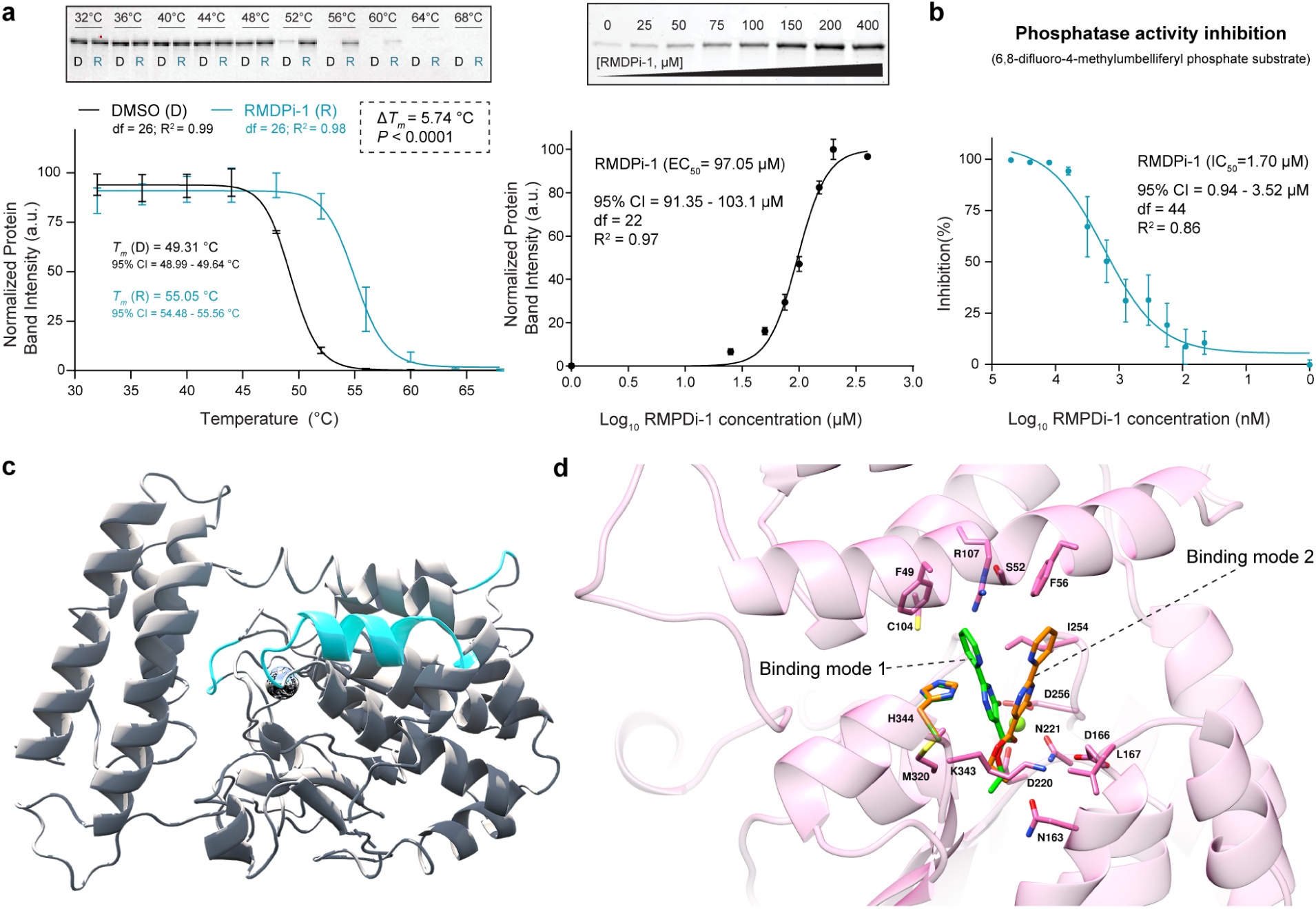
AtRMDP1 Phosphatase activity is inhibited by RMDPi-1 binding to its active site. **a** (left) RMDPi-1 increased the thermal stability AtRMDP1 protein (At2g17430) *in vitro*. Recombinant AtRMDP1 was purified and incubated at 2 µM with 0.5 mM CoCl2 in the presence of either 1% DMSO (D) or 200 µM RMDPi-1 (R). Aliquots were subjected to a temperature gradient ranging from 32°C to 68°C. RMDPi-1 treatment increased the melting temperature (*Tm*) by 5.74°C compared to the DMSO control. Data were fit to a Sigmoidal 4PL model; *F* [1, 52] = 285.9, *P* < 0.0001. Data are plotted as mean ± SE for **a, b**. (right) Thermal stability of AtRMDP1 at 55°C showed a dose-dependent response to RMDPi-1. **b** AtRMDP1 phosphatase activity is inhibited by RMDPi-1. Enzymatic activity (0.24 µM AtRMDP1) was measured using the EnzChek® Phosphatase Assay Kit (Invitrogen™) in the presence of increasing concentrations of RMDPi-1, indicating an IC₅₀ of 1.70 µM. **c** Crystal structure of AtRMDP1 resolved at 1.69Å. The structural analysis revealed residues that were missing in the previously published structure by Bitto et al. (2005), 1XFI (marked in blue and compared in Supplementary file S5). **d** Modelling of the AtRMDP1/ RMDPi-1 binding modes based on the electron densities obtained from the ligand-protein complex structure.

Hit-to-lead optimisation through a virtual screening campaign identified additional analog compounds (Supplementary file S1). Among them was analog 16 (RMDPi-1A16), which had the second-highest predicted binding score and was one of several analogs (along with RMDPi-1) that effectively generated protein crystals via co-incubation with AtRMDP1. Protein crystallography of AtRMDP1 in the presence of RMDPi-1A16 generated at 1.69Å resolution structure containing residues missing from the previously published structure of AtRMDP1 in Bitto et al. (2005) (Figure 2c, Supplementary file S5). The longest portion of the newly resolved peptide structure is a flexible alpha helix neighbouring the active site (Figure 2c). An additional structure resolved at 2.1Å revealed ligand-protein complexes. Modelling of RMDPi-1 in AtRMDP1 showed two alternative binding modes. In both poses, the ligand is anchored through bidentate chelation of the active-site Mg^2^+ ion (or possibly other divalent cations), involving the carbonyl oxygen atoms of the dihydropyrimidinone core and the pendant ester group. The two modes differ principally in the orientation of the ligand scaffold and the positioning of its distal pyridine ring within the surrounding pocket. In both predicted orientations, the pyridine ring extends into a channel delineated by F49, F56, C104, M320, H344, and I254. One binding mode (green coloured, Figure 2d) places the pyridine ring in an orientation compatible with an edge-to-face aromatic interaction with F49, whereas the alternative mode (orange coloured, Figure 2d) adopts a shifted arrangement that preserves metal chelation while engaging hydrophobic/aromatic contacts with F56. The polar side chains of S52 and R107 line the rim of this upper cavity, indicating a possible vector for the introduction of polar substituents in future optimisation (Figure 2d).

Together, these data establish RMDPi-1 as a competitive inhibitor of AtRMDP1 and, through a newly resolved co-crystal structure, define a binding mode in which the ligand chelates the active-site metal ion, blocking phosphometabolite substrate access.

### Loss of RMDP promotes autophagy activation

We assessed the impact of RMDP phosphatase activity on autophagy in representatives of chlorophyte and streptophyte lineages - the unicellular green alga Chlamydomonas and the land plant Arabidopsis. Autophagic flux was assessed in Chlamydomonas strains expressing mVenus-ATG8 in control and *Crrmdp* knockout backgrounds using a canonical GFP-cleavage assay (Shin et al. 2014). The mVenus fluorescent protein is a GFP variant that, like GFP, resists vacuolar proteolysis, so cleavage of an ATG8 fusion protein releases stable free mVenus as a readout of the influx of autophagosomes to the vacuole (Zou et al. 2026). In the control background, RMDPi-1 treatment induced autophagy relative to the DMSO control, and loss of *CrRMDP* similarly elevated basal autophagic flux. Treatment with RMDPi-1 did not further increase autophagic turnover in *CrRMDP* mutants (Figure 3 a).

**Figure 3.**
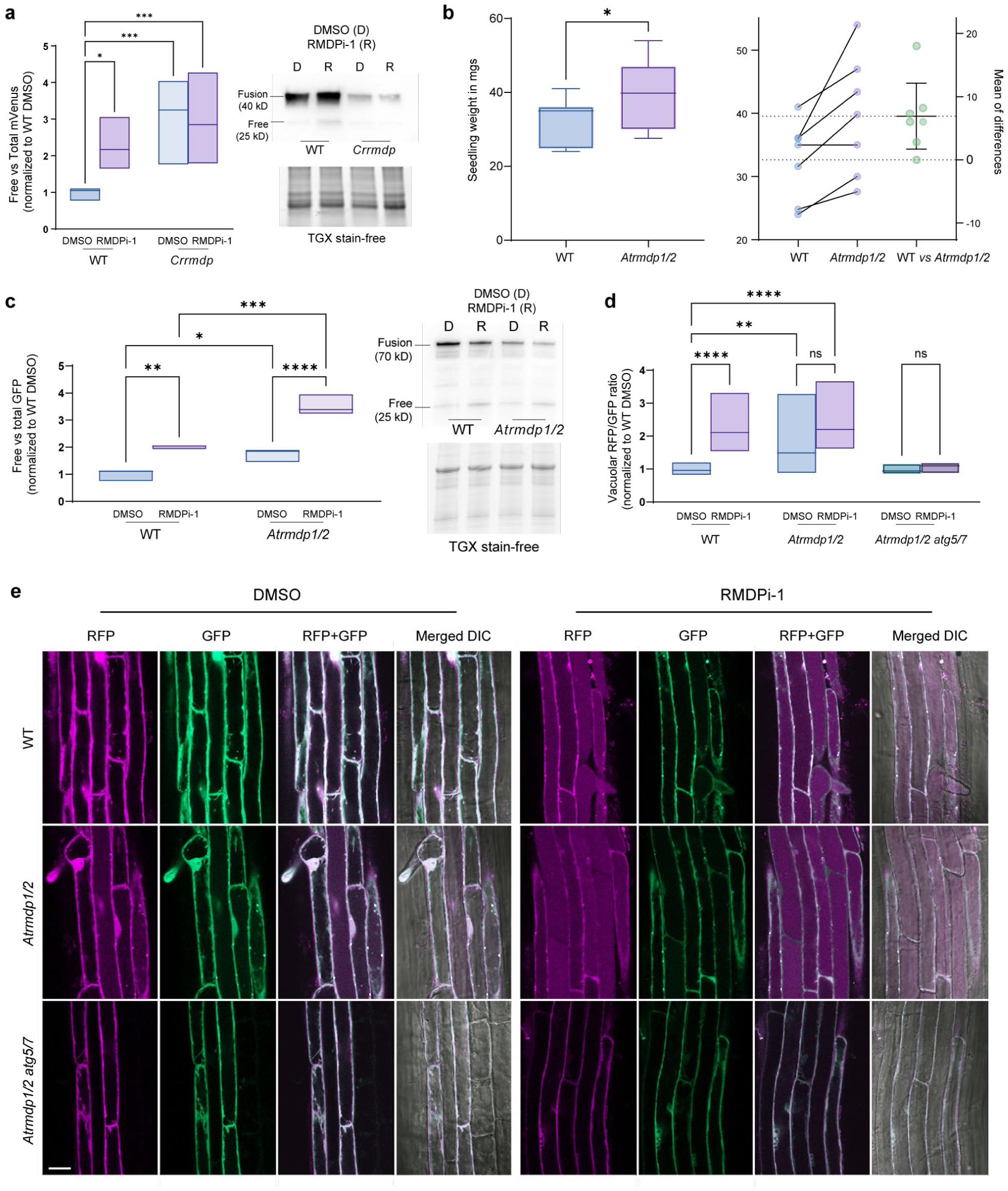
Loss of RMDP enhances autophagy in Arabidopsis and Chlamydomonas. **a** RMDPi-1 treatment and *CrRMDP* knockout induce autophagy in Chlamydomonas. mVenus-cleavage assay using an mVenus-ATG8 reporter line in a WT background following treatment with 50 µM RMDPi-1 for 4 h. Two-way ANOVA, Tukey test, n = 6 biological replicates. **b** Biomass analysis of 13-day-old Arabidopsis seedlings shows increased seedling weight in *rmdp1/2/3* compared to WT grown under identical conditions. Paired Student’s t-test, n = 7 biological replicates. **c** GFP-cleavage assays for Arabidopsis seedlings expressing the pHusion marker (RFP-GFP-ATG8a) in WT and *rmdp1/2* with *rmdp1/2* and RMDPi-1 treatment showing a cumulative effect on autophagic activity. Two-way ANOVA, Tukey test, n = 3 biological replicates. For experiments in **c, d,** and **e**, a 20 µM RMDPi-1 concentration was applied for 6 hours before sampling. **d, e** Tandem tag assay for Arabidopsis seedlings with WT, *rmdp1/2* and *rmdp1/2 atg5/7* (autophagy-deficient control) confirmed the GFP-cleavage assay results. Both RMDPi-1 treatment and AtRMDP1/2 loss result in increased autophagic flux. Two-way ANOVA, Tukey test, n ≥ 9 biological replicates; letters represent significantly different means. Scale bar: 20 µm. For statistical tests, * *P* < 0.05; ** *P* < 0.01; *** *P* < 0.005; **** *P* < 0.001.

In Arabidopsis, three AtRMDP paralogs, AtRMDP1, AtRMDP2 and AtRMDP3 and a pantothenate kinase-RMDP fusion protein, AtPANK2, are present. Single *Atrmdp* knockout lines, generated through T-DNA insertion, did not display detectable phenotypic differences in seedling growth, plant height, or seed yield relative to WT plants (Supplementary file S6). To address potential functional redundancy, we generated double *Atrmdp1/2* and triple *Atrmdp1/2/3* knockout lines using CRISPR-Cas9-mediated genome editing in WT and *Atrmdp3* (T-DNA line) deficient background. (Lee et al. 2018). Triple knockout seedlings exhibited a moderate increase in whole-seedling biomass 13 days after germination (Figure 3b). We established *Atrmdp1/2* seedlings expressing the pHusion-ATG8a reporter (Guichard et al. 2023) to perform a GFP-cleavage assay analogous to that used in Chlamydomonas. Similarly, RMDPi-1 treatment and *AtRMDP1/2* knockout increased free GFP levels, indicating elevated autophagic flux relative to WT controls. In contrast to Chlamydomonas, RMDPi-1 treatment of *Atrmdp1/2* seedlings resulted in a further increase in GFP cleavage, indicating an additive effect of chemical treatment with the loss of two out of four DUF89 family genes on autophagic activity (Figure 3c). The increase in autophagic flux was validated by the TT assay in epidermal root cells of *Atrmdp1/2* seedlings (Figure 3d, e). Irrespective of redundancy, we concluded that AtRMDPs modulate autophagy.

Taken together, genetic loss of *RMDPs* is sufficient to promote autophagy in two distantly related plant lineages, phenocopying RMDPi-1 treatment and supporting RMDPs as the target through which RMDPi-1 enhances autophagic flux.

### RMDP suppression elevates ROS and alters oxidative stress tolerance

RMDP proteins are believed to mitigate cellular damage based on *in vitro* data demonstrating their ability to dephosphorylate phosphate sugars, including potent glycating agents and damaged intermediates of CoA biosynthesis (Huang et al. 2016). Furthermore, HsPANK4 RMDP activity plays an important role in regulating cellular ROS levels (Vella et al. 2024). Therefore, we investigated whether RMDPi-1-mediated inhibition of AtRMDPs affects ROS levels using Arabidopsis roGFP-Orp1 reporter lines that detect hydrogen peroxide accumulation (Nietzel et al. 2019). Treatment with RMDPi-1 resulted in increased ROS levels (Figure 4a). To further investigate oxidative stress responses in the absence of AtRMDPs, superoxide accumulation was visualized by dihydroethidium (DHE) staining in *Atrmdp1/2/3* seedlings. Under control conditions, *Atrmdp1/2/3* seedlings exhibited elevated basal superoxide levels compared to the wild type. Furthermore, treatment with either RMDPi-1 or exposure to methyl viologen (MV), a known inducer of superoxide production, did not further increase superoxide accumulation in the mutant background (Figure 4b, c).

**Figure 4.**
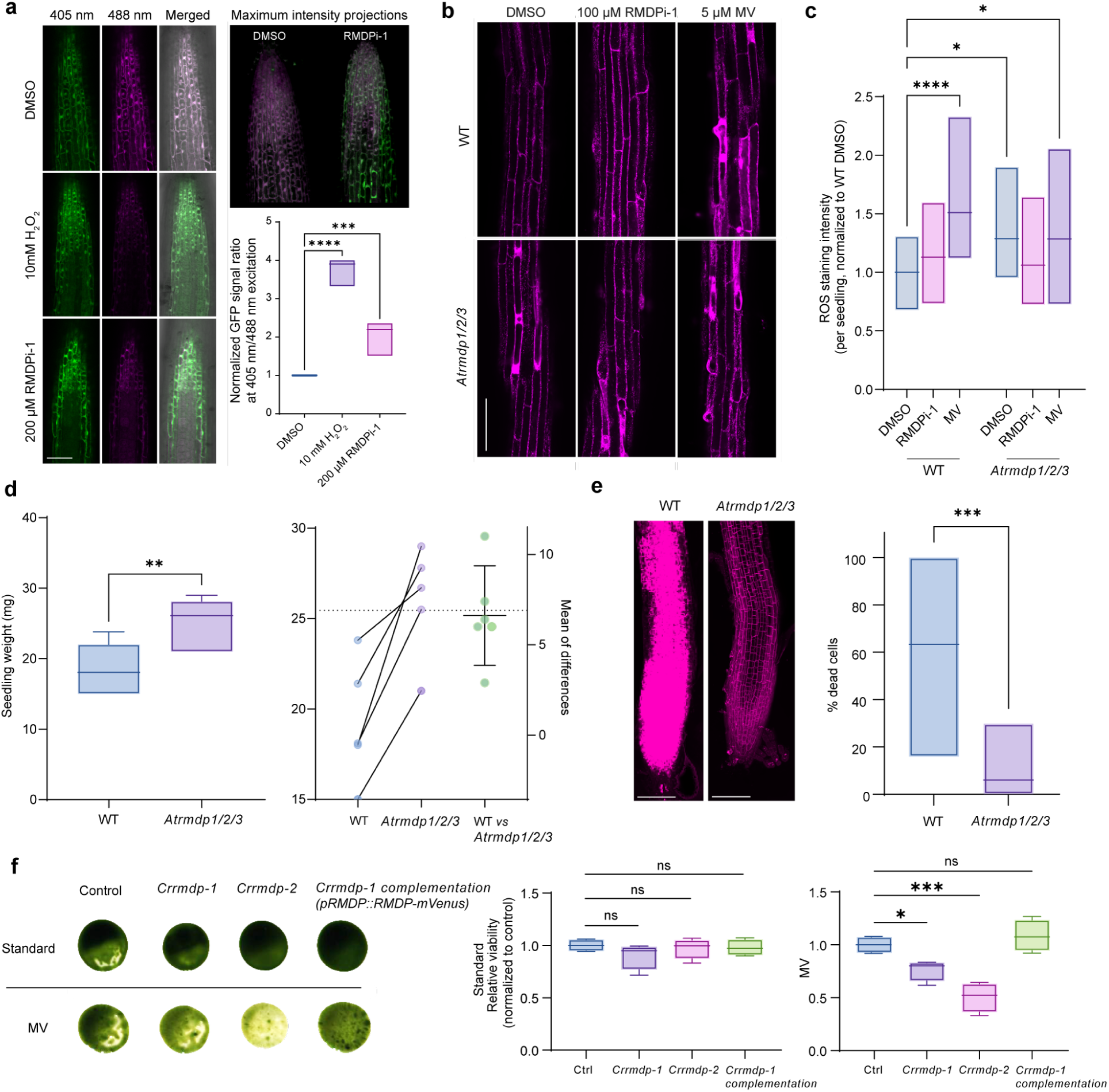
Loss of RMDP activity elevates ROS and alters oxidative stress tolerance. **a** ROS visualization in 6-day-old WT Arabidopsis seedlings with roGFP-Orp1 reporter. Short-term treatment with 200 µM RMDPi-1 increased ROS. One-way ANOVA, Dunnet’s test, n ≥ 4 biological replicates, 50 µm scale bar. **b** ROS staining with DHE in 6-day-old WT and *rmdp1/2/3* Arabidopsis seedlings treated with 100 µM RMDPi-1 or 5 µM MV indicates an increase in ROS in *rmdp1/2/3* seedlings. 50 µM scale bar **c** Quantification of b, one-way ANOVA, Dunnet’s test, n ≥ 9 biological replicates. **d** Biomass analysis of 13-day-old Arabidopsis seedlings transferred to 0.1 µM MV plates showed increased seedling weight in *rmdp1/2/3* compared to WT grown under identical conditions. Paired Student’s t-test, n = 6 biological replicates. **e** Confocal micrographs of roots from WT and *rmdp1/2/3* growing under the same conditions. Dead cells were stained with PI and quantified. Loss of *rmdp1*/2/3 seedlings showed higher MV resistance. Student’s t-test, n = 10 biological replicates. **f** Chlamydomonas growth assay after 10 days for control, two *rmdp* mutants, and an *RMDP* complementation line on standard and 0.4 µM MV-supplemented media, shows decreased MV resistance of mutant lines. Backgrounds were removed using Adobe Photoshop CC; adjustments to brightness and contrast were applied uniformly. One-way ANOVA, Dunnet’s test, n = 4 biological replicates. \**P* < 0.05; \*\**P* < 0.01, \*\*\**P* < 0.0005, **** *P* < 0.0001.

To assess whether elevated basal ROS levels influenced oxidative stress tolerance, we evaluated the response of *Atrmdp1/2/3* seedlings to MV. Six-day-old seedlings were transferred to MV-containing medium and grown for an additional week. Under these conditions, *Atrmdp1/2/3* seedlings had greater overall biomass than WT controls, indicating enhanced resistance to MV-induced stress (Figure 4d). To quantify cellular damage, we stained root tips with propidium iodide (PI) and quantified dead cells in the meristematic and elongation zones. The *Atrmdp1/2/3* roots had a significantly lower number of dead cells per root compared to WT seedlings following MV treatment (Figure 4e), consistent with increased oxidative stress tolerance. Conversely, in Chlamydomonas, *rmdp* mutants grown on MV-supplemented medium showed reduced growth relative to the wild-type and a complemented line (Figure 4f), indicating increased rather than decreased sensitivity to oxidative stress. Furthermore, Chlamydomonas *rmdp* knockout mutants showed a growth phenotype indistinguishable from the control (Figure 4f). Loss of RMDP activity therefore has opposing consequences in the two organisms. Arabidopsis *rmdp1/2/3* mutants display elevated basal ROS yet enhanced tolerance to acute oxidative stress in non-photosynthetic roots, whereas photosynthetic Chlamydomonas *rmdp* mutants are more sensitive.

### RMDP loss drives sugar-phosphate accumulation and redox imbalance

To understand the physiological consequences of RMDP loss, we profiled metabolomic changes in Arabidopsis and Chlamydomonas. We predicted affected metabolites based on the distinct *in vitro* substrate preferences of standalone RMDP versus fusion-type RMDP enzymes. AtRMDP1 acts preferentially on sugar phosphates, particularly hexose, pentose, and tetrose phosphates (Huang et al. 2016). In contrast, the fusion proteins AtPANK2 and HsPANK4 act preferentially on CoA-related metabolites (Huang et al. 2016), which is consistent with PANK’s central role in CoA synthesis, where it salvages reactive intermediates. CrRMDP, CrPANK and their substrate preferences have not been previously characterized.

As anticipated, the levels of detected masses consistent with an unidentified hexose phosphate (UHP1) rose in Arabidopsis seedlings following the loss of AtRMDP1/2/3 and RMDPi-1 treatment (Figure 5a). The level of an unidentified pentose phosphate (UPP1) increased significantly only when knockout and RMDPi-1 treatment were combined (Figure 5a). This additive effect on UPP1 can be explained because AtPANK2, which remains intact in the *rmdp1/2/3* background, is predicted to bind RMDPi-1 and act weakly on phosphate sugars, in agreement with previous *in vitro* studies (Huang et al. 2016). RMDPi-1 inhibition could further increase UPP1. Furthermore, 4’-phosphopantothenate - the first committed intermediate of CoA synthesis and itself a product of AtPANK2 - was also detected. RMDPi-1 treatment of WT plants raised 4’-phosphopantothenate more than the *rmdp1/2/3* knockout alone, consistent with inhibition of AtPANK2. Combining the *rmdp1/2/3* knockout with RMDPi-1 treatment produced a further, additive increase (Figure 5a). In Chlamydomonas *rmdp* knockouts, phosphorylated sugars accumulated strongly, most notably erythrose-4-phosphate, glucose 6-phosphate, fructose 6-phosphate and two unknown UPPs (Figure 5b).

**Figure 5.**
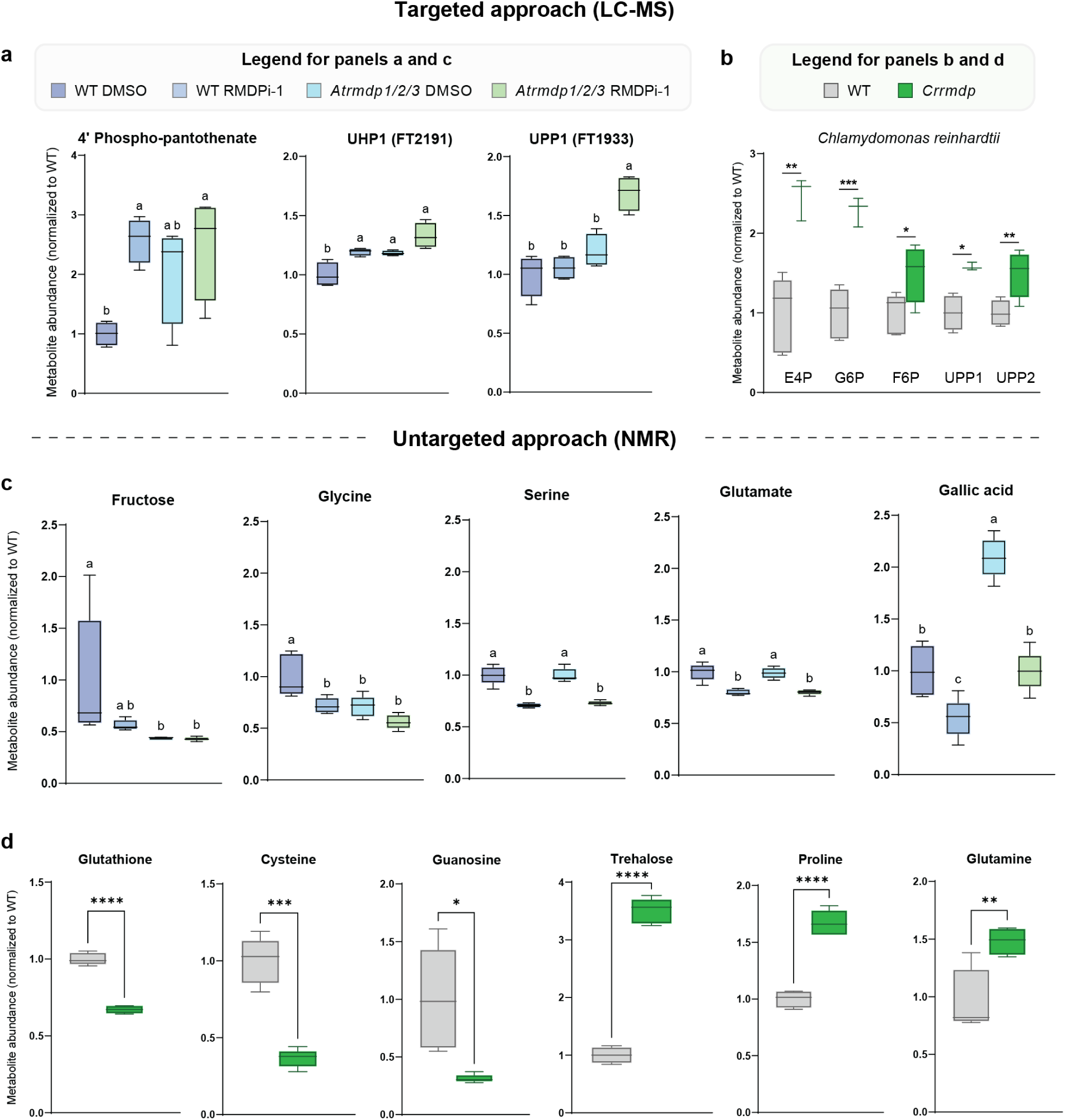
Select metabolites detected via targeted and untargeted metabolomic approaches. **a** Masses corresponding to phosphate metabolites enriched following RMDPi-1 or knockdown of *rmdp1/2/3* in Arabidopsis seedlings include 4’-phosphopantothenate, Unknown Hexose Phosphate 1 (UHP1) and Unknown Pentose Phosphate 1 (UPP1). **b** In Chlamydomonas, masses corresponding to erythose-4-phosphate (E4P) were enriched in a *rmdp* knockdown, along with glucose 6-phosphate (G6P), fructose 6-phosphate (F6P) and UPP1/2. Untargeted NMR-based metabolomics revealed differentially enriched metabolites in Arabidopsis (**c**) and Chlamydomonas (**d**). All data were normalised to the WT vehicle-treated control and are presented as Tukey boxplots, n ≥ 3 biological replicates. For a, c, One-way ANOVA with Tukey test, letters indicate means with significant differences; b, d, Welch’s t-test (two-tailed), * *P* < 0.05; ** *P* < 0.01; *** *P* < 0.005; **** *P* < 0.001. For the complete list of detected metabolites, see supplementary file S7.

An untargeted NMR approach identified additional metabolites suggesting reduced hexose dephosphorylation, with lower fructose levels in *rmdp1/2/3* seedlings and following RMDPi-1 treatment. Glycine showed a similar pattern of inhibition in *rmdp1/2/3* seedlings with and without RMDPi-1, whereas serine and glutamate were reduced only after RMDPi-1 treatment (Figure 5c). Glutathione, a tripeptide of glutamate, cysteine, and glycine, is a principal cellular ROS scavenger. We also detected the antioxidant gallic acid in Arabidopsis samples. The RMDPi-1 treatments lowered abundance, whereas the *rmdp1/2/3* control seedlings had significantly higher amounts. In Chlamydomonas *rmdp* mutants, glutathione and cysteine were both depleted, along with guanosine. In parallel, the osmoprotectants proline, trehalose and glutamine accumulated, indicating a broader mobilization of protective redox metabolites, resembling the Arabidopsis response (Figure 5d). Metabolomic profiling of Arabidopsis and Chlamydomonas supports that loss of RMDP activity drives accumulation of sugar phosphates and 4’phospho-pantothenate, alongside a selective depletion of ROS-protective metabolites. For the complete list of detected metabolites, see Supplementary file S7.

## Discussion

Chemical genetics approaches have long served as powerful entry points into unexplored biological pathways. Here, we applied this principle to RMDPi-1, a novel small-molecule enhancer of autophagy in Arabidopsis. Integrating chemical genetics and an *in vivo* PAL workflow, we identified AtRMDPs as targets of RMDPi-1. Significantly, these phosphatases have no prior connection to ROS-mediated autophagy signalling or stress resilience. We provide multiple independent lines of evidence that RMDPs have conserved metabolite damage control functions and that their inhibition stimulates ROS-mediated autophagy.

Our conclusion that AtRMDPs are the functionally relevant targets of RMDPi-1 is supported by *in vitro* biophysical characterisation confirming competitive inhibition of AtRMDP1 by RMDPi-1, and crystallisation of AtRMDP1 with RMDPi-1 that enabled modelling of the binding modes of RMDPi-1 within the catalytic site (Figure 2d). The structural data provides a high-resolution blueprint for improvement of compound potency and selectivity. As with any small organic molecule, off-target activity cannot be excluded, and future work with more selective analogs guided by the structural data reported here will be important to delineate the contribution of such interactions to the observed phenotypes.

The effect of RMDPi-1 on autophagy is conserved across the chlorophyte and streptophyte lineages. In Chlamydomonas, knockout of CrRMDP confers resistance to RMDPi-1, directly linking loss of the target to compound insensitivity (Figure 3 a). In Arabidopsis, treating the *rmdp1/2* knockout with RMDPi-1 enhanced autophagy synergistically, an effect seen by western blot but not by the tandem-tag assay. Because RMDPi-1 still increased flux beyond the knockout alone, the compound must also act on the remaining paralog, RMDP3. This confirms that the AtRMDPs are the relevant targets and shows that the paralogs are partly redundant. Importantly, pharmacological inhibition and genetic knockout converge on a similarly enhanced autophagic flux in both organisms (Figure 3c). Together, these results establish the RMDPs as bona fide targets for stimulating autophagic flux in plants.

The link we identified between AtRMDP proteins and autophagy regulation has not previously been proposed, but indirect evidence exists. A recently developed PUPylation-based proximity labelling assay in Arabidopsis shows that TOR physically associates with AtPANK2 (Zheng et al. 2025), directly coupling the PANK-RMDP fusion protein to the master autophagy-regulatory kinase. A parallel for such regulation comes from mammalian CD4^+^ T cells, where PDK1 binds PANK4 directly and phosphorylates it, inhibiting its phosphatase activity and thereby de-repressing CoA synthesis to support *de novo* lipid synthesis and proliferation (Hwang et al. 2025). Whether this mechanism reflects TOR-dependent phosphorylation of AtPANK2, a reciprocal influence of CoA pathway status on TOR activity, or both, warrants further investigation. Furthermore, a genome-wide deletion screen in *Schizosaccharomyces pombe* identified the AtRMDP homolog, YMR027W, among mutants with mild sensitivity to rapamycin (Doi et al. 2015), a canonical TORC1 inhibitor and autophagy inducer; however, this observation lacked mechanistic context. Our finding that *AtRMDP* knockouts themselves promote autophagy responses in plant cells now provides a possible explanation. Rapamycin imposes a second, independent autophagy-inducing signal on YMR027W-deleted cells already experiencing elevated autophagic flux, driving it past a cytoprotective threshold and explaining the lethality (Liu & Levine 2015; Kriel & Loos 2019). These observations extend the functional connection between *RMDP* loss and autophagy induction further to fungi.

How does AtRMDP inhibition induce autophagy? The existing literature supports two mechanistic routes, which are not mutually exclusive. Arabidopsis encodes three AtRMDP proteins and a bifunctional AtPANK2 carrying both a pantothenate kinase and a RMDP domain with conserved active sites (Huang et al. 2016). AtRMDP1 shows high *in vitro* activity against sugar phosphates, particularly ribose 5-phosphate and erythrose 4-phosphate, that are among the most potent endogenous glycating agents, with only modest activity against CoA biosynthesis-related substrates. AtPANK2 and its human orthologue HsPANK4, by contrast, preferentially dephosphorylate the acetyl-CoA precursors: 4’-phosphopantothenate (the first committed step in CoA biosynthesis), 4’-phosphopantetheine, and its oxidized derivative S-sulfonate, with limited activity against sugar phosphates (Huang et al. 2016). This substrate divergence has direct mechanistic implications.

In the first route, AtPANK2 perturbation may alter cellular CoA levels. HsPANK4 hydrolyses 4’-phosphopantothenate, 4’-phosphopantetheine and its oxidized derivatives at rates sufficient to influence CoA pathway flux, functioning simultaneously as a repair enzyme and a regulator of cellular CoA levels (Dennis et al. 2020; Dibble et al. 2022). Muscle-specific HsPank4 deletion in mice elevates intramuscular acetyl-CoA, demonstrating that its phosphatase activity has measurable metabolic consequences *in vivo* (Miranda-Cervantes et al. 2025). Our metabolomic data support this, with both RMDPi-1 and *rmdp1/2/3* raising 4’-phosphopantothenate levels in Arabidopsis (Figure 5a). Given that cytosolic acetyl-CoA directly suppresses autophagy via mTORC1 (Mariño et al. 2014), reduced CoA levels resulting from AtPANK2 perturbation represent a plausible route through which RMDP dysfunction could enhance autophagic flux in both plants and mammals.

In the second route, RMDPi-1-mediated inhibition of RMDP disrupts glycation damage control and central carbon metabolism, allowing reactive sugar phosphates to accumulate. Interestingly, in Arabidopsis, we detected a significant drop in masses corresponding to fructose, a known signal for SnRK1 activation, which inhibits TOR, and thereby induces autophagy (Janse van Rensburg et al. 2019). Huang et al. established that DUF89 proteins function as metabolite damage-control enzymes whose primary role is the pre-emptive removal of aberrant phosphometabolites. Metabolite accumulation can cause macromolecular damage and elevated intracellular ROS levels; two established triggers of autophagy. We observed that a loss of RMDP causes phosphate sugars to accumulate in both Arabidopsis and Chlamydomonas (Figure 5a-b). Further evidence from human cancer biology independently reinforces this connection. In glioblastoma, HsPANK4 is induced by temozolomide treatment, and its expression correlates with acquired drug resistance. HsPANK4 depletion restores drug sensitivity through intracellular ROS accumulation and cell death, phenotypes that are strictly dependent on an intact RMDP domain and are accompanied by broad downregulation of ROS-scavenging and detoxification proteins (Vella et al. 2024). Our metabolomic data support this model.

Arabidopsis *rmdp* mutants accumulated the antioxidant gallic acid, and Chlamydomonas *rmdp* mutants accumulated the ROS-scavengers trehalose (Luo et al. 2008) and proline (Rehman et al. 2021), whereas cysteine and glutathione (Gill & Tuteja 2010) were depleted (Figure 5d), consistent with sustained oxidative demand, whereby the elevated ROS drives glutathione consumption faster than it can be regenerated, potentially drawing down its precursors. The accumulation of proline and trehalose, osmoprotectants with ROS-scavenging roles, fits a broader protective response. Together, these findings underscore the importance of RMDP-mediated metabolite repair for cellular redox homeostasis, since loss of this capacity sensitizes cells to specific stress challenges. Comparable associations recur across other stress contexts in plants. An RMDP ortholog is upregulated under salt stress in the halophyte *Schrenkiellaparvula* (§ekerci et al. 2024), after high-temperature stress in *Sorbus pohuashanensis* (Pei et al. 2022), and AtRMDP2 is enriched in Arabidopsis upon iron deficiency, alongside ROS scavengers (Lan et al. 2011).

RMDPi-1 treatment and *AtRMDP* knockout stimulate ROS accumulation in Arabidopsis. Yet rather than causing toxicity, this effect is accompanied by improved stress adaptation. Triple knockout *rmdp1/2/3* plants display higher basal ROS levels, elevated basal autophagy, and markedly greater resistance to MV (Figure 4a-d), a herbicide whose toxicity derives from superoxide anion formation, as evidenced by reduced cytotoxicity in root cells and enhanced seedling growth relative to wild type. This result is consistent with the established concept that moderate, controlled ROS acts as an adaptive signal in plants, priming stress tolerance responses in a hormetic manner (Agathokleous et al. 2019; Mittler et al. 2022). The magnitude of ROS elevation therefore appears to determine whether the outcome is adaptive or damaging, a balance likely shaped by genetic redundancy and cellular context. In Arabidopsis, loss of *AtRMDP* genes alone may not fully abolish glycation damage control, as residual AtPANK2 activity may partially compensate, keeping ROS within a beneficial range. By contrast, knockout of the single *RMDP* gene in Chlamydomonas had a more pronounced effect despite also encoding a PANK2-RMDP fusion protein. Because chloroplast electron transport is a major ROS source and MV generates superoxide by accepting electrons from photosystem I, oxidative damage is expected to be more acute in photosynthetic Chlamydomonas cells than in the largely non-photosynthetic Arabidopsis root tissue examined here. Thus, the heightened MV sensitivity of *rmdp* in Chlamydomonas is predictable. More broadly, perturbation of metabolite damage-control systems typically produces subtle fitness costs that are not apparent under standard conditions but emerge under stress, precisely the pattern observed here in AtRMDP loss-of-function lines.

Our results demonstrate the physiological impacts of RMDP suppression and establish RMDP/DUF89 phosphatases as targets to influence ROS and autophagy signaling through their conserved reactive metabolite damage-control functions, while positioning RMDPi-1 as a novel chemical tool to modulate autophagic flux across evolutionarily distant systems.

## Materials and Methods

### Plant materials

*Arabidopsis thaliana* seeds were surface sterilised with bleach solution (2.7 g/l sodium hypochlorite, 0.05% (v/v) Tween 20) for 30 to 40 min, washed three times with sterile Milli-Q water and plated on Murashige and Skoog plates (0.5x MS complete Murashige and Skoog, 10 mM 2-(N-morpholino)ethanesulfonic acid (MES), 1% (w/v) Sucrose, 0.8% (w/v) plant agar, pH 5.8). The seeds were vernalised on plates at 4°C in darkness for 24h before they were placed vertically in long light conditions (16 h of 150 µM photons m^2^ s^-1^ light at 22°C, 8 h of dark at 20°C). For Arabidopsis plants cultivated in pots, S-Jord soil (Hasselfors, Sweden) was used, and grown in long-day conditions (16 h of 150 µM photons m^2^ s^-1^ light at 22°C, 8 h of dark at 20°C and 70% relative humidity). The Arabidopsis lines utilised in this study included Wild-type (Columbia-0 ecotype) *Arabidopsis thaliana* (L.) Heynh (taxonomy ID: 3702); TT-ATG8a, which consists of a tandem tag of TagRFP and mWasabi fused to ATG8 (2x35S::TagRFP-mWasabi-ATG8a; Dauphinee et al. 2019); roGFP-orp1 (Nietzel et al. 2019) and the T-DNA insertion line DUF89 - At2g17340.1 knock-out (SALK_143093C), obtained from the Nottingham Arabidopsis Stock Centre (NASC).

### Tandem Tag assay

The assay was performed as described in Dauphinee et al. (2020) on TagRFP-mWasabi-ATG8a Arabidopsis marker lines or pHusion (Guichard et al. 2023) tandem tag marker lines. AZD8055 (Dong et al. 2015) served as a positive control for autophagy induction.

### Chemicals, synthesis and photoaffinity labelling pulldown assay

Endosidin 18 (CAS# 60060-10-8) was purchased from Maybridge (SPB05487). Synthesis of specific analogues, probe molecules and biotin was performed through a collaboration with the Chemical Biology Consortium of Sweden (CBCS). See supplementary file S2 for details.

#### Photoaffinity labelling

One-week-old seedlings were transferred to a 6-well plate containing 3 ml Murashige and Skoog medium (0.5x MS complete Murashige and Skoog, 10 mM 2-(N-morpholino)ethanesulfonic acid (MES), 1% (w/v) Sucrose, pH 5.8). The seedlings were fully submerged and treated with DMSO, RMDPi-1-probe and the control probe. DMSO was added to each treatment to a final concentration of 1% (v/v) DMSO. After incubating in the dark at 22°C for 6 hours, seedlings were rinsed three times with 500 µl liquid Murashige and Skoog medium. To prevent dehydration of the seedlings, 100 µl Murashige and Skoog medium per well was retained. The seedlings were evenly distributed on the bottom of each well to maintain a consistent 3 cm distance from the UV source during photo-crosslinking, which occurred for 20 minutes on the tabletop using an 8-watt UV lamp at 365 nm (Thermo Scientific, PI95035).

#### Protein extraction

Whole seedlings were frozen in liquid nitrogen and homogenised with a mortar and pestle. The plant material was transferred to 1.5 ml low protein binding tubes (ThermoFisher Scientific™). PAL extraction buffer (50 mM Tris HCl, 150 mM NaCl, 0.1% SDS, pH 7.5) supplemented with 1xComplete protease inhibitor (EDTA free, Roche) was added (5 µl per mg seedling), and the plant material was thawed on ice. After vortexing and centrifugation (13,300 rpm, 15 minutes, 4°C), protein concentration of the supernatant was determined using the Qubit protein assay kit as described in Mol. Probes Life Technology, 2010.

#### Cyclo-addition of biotin-azide tag

A freshly prepared 10x Click Mix (0.5 volumes of 50 mM CuSO4, 0.5 volumes of 50 mM tris(2-carboxyethyl)phosphine (TCEP) pH 7.0, and 1.5 volumes of 2 mM tris[(1-benzyl-1H-1,2,3-triazol-4-yl)methyl]amine (TBTA) in 80% (v/v) tert-Butanol and 20% (v/v) DMSO) was used. For the no-click control, CuSO4 was substituted for Milli-Q water. The click-mix was thoroughly vortexed before use. 0.5 mM biotin-azide was added in a final concentration of 25 µM. After a 30-minute incubation at 32°C, the reaction was stopped by adding 5 mM EDTA. EDTA was only added to click reaction samples. Excess biotin-azide was removed using Zeba columns (ThermoFisher Scientific™). The samples were loaded to the top of the resin and 15 µl Milli-Q water was added to the column and centrifuged at 1500xg for 2 minutes at 4°C.

#### Purification with Streptavidin-magnetic beads

The desalted protein sample was diluted to 150 µl with wash buffer (50 mM Tris HCl, 25 mM NaCl pH 7.5) supplemented with 1x Complete protease inhibitor (EDTA free, Roche). Streptavidin-coated magnetic beads (Sera-Mag SpeedBeads, GE Life Sciences) were prepared by mixing 20 µl with 10x the volume of the wash buffer. The tube was inverted several times, and the beads were captured using a MagneSphere Magnetic Separation Stand (Promega). The beads were resuspended in 25 µl wash buffer per sample. 25 µl of beads were added to each sample and the samples were incubated for 2 hours at 4°C with mixing by inversion. After incubation, the beads were captured on the magnetic stand and washed three times with 1 ml wash buffer. Afterwards, the beads were resuspended in 100 µl wash buffer for LC-MS analysis, or in 20 µl 1x Laemmli buffer (Bio-Rad) with 5% (v/v) β-mercaptoethanol (AppliChem) and heated up for 5 minutes at 65°C for testing the PAL pulldown assay with SDS-PAGE on a Bio-Rad Criterion 12.5% SDS-PAGE gel. The SDS-PAGE samples were spun down at 13,000 rpm for 5 minutes at 4°C. The SDS-PAGE gels were stained with Oriole™ stain (Bio-Rad). The gels were incubated in 50 ml of 1x stain solution and rocked vigorously for 90 minutes and imaged on the GelDoc XR (Bio-Rad) at 270 nm /604 nm.

### Structure validation and purity assessment of RMDPi-1

RMDPi-1 was dissolved in DMSO-d6 with 0.05% TMS to obtain a 20 mM solution. 1H and 13C NMR spectra were recorded at 25°C on a Bruker Avance III 600 MHz spectrometer equipped with a 5 mm 1H/13C/15N/31P inverse detection CryoProbe. 2D 1H,1H-TOCSY, 1H,1H-NOESY, 1H,13C-HSQC, and 1H,13C-HMBC spectra were recorded to confirm the structure. NMR spectra were processed with TopSpin 4.0.6 (Bruker) and chemical shifts were calibrated using the TMS peak: 5H 0.00 ppm and 5C 0.00 ppm. LC-MS: MS analysis was carried out on an Orbitrap QExactive mass spectrometer (ThermoFisher Scientific™) with chromatographic separation performed on a Vanquish Horizon UHPLC system (ThermoFisher Scientific™). The UHPLC system was equipped with a C18 column (2.1*100 mm) using a mobile phase of water and acetonitrile (linear gradient) and a flow rate of 0.4 ml/min. RMDPi-1 was dissolved in methanol from which 0.5 µl was injected.

1H NMR (DMSO-d6): 5 12.71 (br s, 1H, NH), 8.79 (ddd, J = 4.8, 1.7, 0.9 Hz, 1H, H6’), 8.54 (br s, 1H, H6), 8.37 (dt, J = 7.9, 1.1 Hz, 1H, H3’), 8.10 (td, J = 7.7, 1.7 Hz, 1H, H4’), 7.71 (ddd, J = 7.7, 4.8, 1.1 Hz, 1H, H5’), 4.25 (q, J = 7.1 Hz, 2H, CH2), 1.29 (t, J = 7.1 Hz, 3H, CH3). 13C NMR (DMSO-d6): 5 163.3 (COO), 157.1 (br, C2), 149.2 (C6’), 147.6 (C2’), 138.0 (C4’), 127.3 (C5’), 123.0 (C3’), 115.9 (br, C5), 60.3 (CH2), 13.9 (CH3). LC-MS: calculated for C12N3O3H12: [M + H]+ m/z 246.09; found m/z 246.09.

Purity estimation based on integrated signals in the 1H NMR spectrum: >99.3% (Supplementary file S7).

### Proteomics

Affinity-enrichment liquid chromatography tandem mass spectrometry (AE-LC-MS/MS) in VIB Proteomics Core, Ghent. Comparison of our modulator RMDPi-1-probe vs control probe. 9 samples were searched together using the MaxQuant algorithm (version 1.6.11.0). False Discovery Rate (FDR) set at 1% on PSM (Peptide-to-Spectrum Matches), peptide and protein level. Spectra were searched against the Arabidopsis reference protein sequences (database release version of April 2020), containing 39,359 sequences (UniProt, taxid3702).

### Western blot and Immunostaining

The proteins were transferred from an SDS-PAGE gel by Bio-Rad turbo blot onto a polyvinylidene difluoride membrane. The membrane was washed with PBST (1 tablet (Medcicago) per litre water, 0.1% Tween 20 (AppliChem)) and blocked with 1% (w/v) dry milk (Merck) in PBST for 15 min at room temperature, followed by three washes for 5 minutes with PBST. The membrane was incubated with 1/5000 diluted Streptavidin-HRP (Horseradish Peroxidase) in 25 ml PBST for 45 min. After a washing step of 3x 5 minutes with PBST, the blots were developed using HRP ECL Prime (Amersham) and GelDoc XR (Bio-Rad).

### AtRMDP1 overexpression

The synthetic full-length coding region of AtRMDP1 (AT2G17340) (Eurofins Genomics) was cloned into the pET-28b(+) vector (Novagen) vector for bacterial expression of N-terminally 6xHis-tagged AtRMDP1 protein. The At2G17340.1 sequence, including point mutations and NcoI and XhoI restriction digestion cleavage sites, is detailed in the supplementary file S8. The Synthetic DNA fragment as well as pET-28b(+) were digested using NcoI and XhoI FastDigest (ThermoFisher Scientific™) and ligated by T4 Ligase (1U, ThermoFisher Scientific™). The ligation product was transformed into *Escherichia coli* Top10 cells and plated on LB plates with 50 µg/ml kanamycin, followed by colony PCR to confirm the presence of AtRMDP1 in the transformed colonies. Primer information in the supplements. For AtRMDP1 overexpression, transformed Rosetta cells were used to inoculate 5 ml of LB media supplemented with antibiotics (ampicillin) and incubated at 37°C overnight. One millilitre of the overnight culture was transferred into 50 ml of LB medium with ampicillin. The cultures were grown at 37°C with shaking at 200 rpm for 3 hours. A 50 ml aliquot of the culture was further inoculated into 450 ml of LB medium with ampicillin. The culture was incubated at 37°C with shaking at 200 rpm. The growth continued until the optical density at 600 nm (OD600) reached approximately 1.0. Overexpression was initiated by adding 50 µl of a 1M IPTG (isopropyl β-D-1-thiogalactopyranoside) solution. Following induction, the incubation temperature was reduced to 20°C, and the culture was shaken at 200 rpm for an additional 20 hours.

### AtRMDP1 protein purification

Rosetta cells with overexpressed AtRMDP1 were resuspended in binding buffer (500 mM NaCl, 35 mM Imidazole, 20 mM sodium phosphate, 20% (v/v) ethylene glycol, 1 mM DTT, pH 7.4) supplemented with 1x Complete protease inhibitor (EDTA free, Roche) and lysed in a cell disruptor at 20 kpsi. The lysed cells were treated with DNase and centrifuged for 20 minutes at 38.000xg at 4°C. The purification of AtRMDP1 from the lysate was conducted using the Akta (Cytiva™) System through immobilised metal ion affinity chromatography (IMAC) and subsequent size exclusion chromatography. HisTrap HP (Cytiva™) columns with insolubilised Ni^2^+ were used to purify his6-tagged AtRMDP1 from the lysate. Elution of the bound AtRMDP1 was achieved using an elution buffer comprising 500 mM NaCl, 350 mM Imidazole, 20 mM sodium phosphate, 1 mM DTT, pH 7.4. Eluted fractions were collected for SDS-PAGE analysis. The eluted fractions were pooled and subjected to size exclusion chromatography using a HiLoad Superdex 200 column (Cytiva™). This step served to further refine the purity of the AtRMDP1 protein and exchange the buffer to HEPES buffer (50 mM HEPES, 100 mM NaCl, pH 7.4). The concentrated and purified AtRMDP1 was flash-frozen in liquid nitrogen and stored at -20°C, preserving its integrity for subsequent experimental analyses.

### Thermal Shift Assay

Purified AtRMDP1 protein in a concentration of 2 µM was supplemented with 0.5 mM CoCl2 as a metal co-factor and mixed at 4°C for 5 minutes. The protein sample was divided into two treatment groups. The first group, serving as a control, was treated with a final concentration of 1% (v/v) DMSO, while the second group was treated with 200 µM RMDPi-1 in 1% (v/v) DMSO. To facilitate drug-protein binding, the samples were rotated in a cold room for 10 minutes. 100 µl of each sample was transferred to individual 0.5 ml PCR tubes in preparation for the subsequent heat-challenge step. The tubes were exposed to a heat gradient from 28°C to 68°C with an increment of 4°C and upon completion, the tubes were promptly chilled on ice for 5 minutes. Protein aggregates were removed by a 40-minute centrifugation at 13.3k rpm at 4°C. The supernatant was carefully collected to avoid disruption of the pellet. The soluble protein faction of the control vs treatment group was visualised by SDS-PAGE on a 15% stain-free TGX gel (BioRad, 26 well) and imaged on GelDoc XR (Bio-Rad). The intensity of the protein bands was quantified with Image Studio™ Light quantification software (Li-COR Biosciences).

### Phosphatase activity assay

The phosphatase activity of purified AtRMDP1 protein was assessed by EnzChek® phosphatase assay kit (Invitrogen™) following the manufacturer’s instructions. For this purpose, a protein concentration of 0.24 µM was pre-incubated with 50 pM CoCl2 for 10 minutes at 4°C. Following the pre-incubation, phosphatase activity was measured in both the 1% DMSO control group and in groups treated with increasing concentrations of RMDPi-1, as well as RMDPi-1-analogs as a negative control. The fluorescence readout was measured by a BMG Labtech POLARstar Omega Microplate Reader.

### Crystallisation, data collection, and processing

At2g17340 was crystallised using the sitting-drop vapour-diffusion method at 20°C. Initial screening was performed with the JCSG+ kit (Page et al. 2003) and Morpheus kit. The protein solution was pre-incubated in ice with RMDPi-1 ligand at 50 µM concentration. Final conditions for crystallization were as follows: 1 ml of protein solution (2.6 mg/ml) was mixed with 1 ml of reservoir solution containing 10% w/v PEG 20 000, 20% v/v PEG MME 550, 0.03 M of each divalent cation, 0.1 M MOPS/HEPES-Na, pH 7.5 (divalent cation: 0.03 M magnesium chloride, 0.03 M calcium chloride). Crystals were flash-cooled in liquid nitrogen. Diffraction data were collected at 100 K at the massif ID30A-1 beam line at the European Synchrotron Radiation Facility (ESRF, Grenoble, France). The data were autoprocessed and scaled using XDS (Kabsch 2010) and aimless. The structure was solved by molecular replacement using Phaser of the CCP4 suite (McCoy et al. 2007). The search model was the previously solved structure of At2g17340 (PDB code 1XFI). The solution was confirmed by the observation of electron density for the residues 173-180 and 271-272, not modelled in the search model. The missing residues in the model were built using ARP/WARP (Beshnova et al. 2017). Rigid body refinement was followed by alternating cycles of restrained refinement in REFMAC5 (Murshudov et al. 1997) and manual rebuilding in O (Jones et al. 1991). Data collection and refinement statistics are presented in Table 1. Atomic coordinates and structure factor data have been deposited at the Protein Data Bank with accession code 32BK.

### Molecular docking

Crystallographic coordinates of Arabidopsis At2g17340 were prepared for docking calculations by removing solvent and buffer molecules and adding hydrogens using REDUCE (v3.23) (Word et al. 1999).

Since no experimental structure of the Chlamydomonas damage-control phosphatase (UniProt A8HSN7) was available, a comparative model was generated with the SWISS-MODEL server (Waterhouse et al. 2018). Chain A of PDB (Berman et al. 2000) entry 2Q40 (Levin et al. 2007), the 1.70 Å refined crystal structure of At2g17340 (UniProt Q949P3, The UniProt Consortium 2025), was selected as template. It shares 42.82% sequence identity with the query, and it covers 80% of the target sequence (modelled range 1-408). The query structure was modelled as a monomer. The final model gave a QMEANDisCo score of 0.66. The Mg^2+^-bound model was used as the receptor for the subsequent structure preparation. The model was energy-minimised with GROMACS 2026.0 (Abraham et al. 2015). The topology was generated using the AMBER99SB-ILDN force field (Lindorff-Larsen et al. 2010) and the TIP3P water model (Jorgensen et al. 1983); the system was solvated in a cubic periodic box and neutralised with Na^+^ or Cl^-^ counterions. Steepest-descent minimisation was run for up to 5000 steps. The minimised structure was used as the receptor for the subsequent docking calculations. The RMDPi-1 ligand was generated by building the structure in Marvin and subsequently protonated to represent the predominant protomer at physiological pH (7.4), using the cxcalc tool (Marvin and cxcalc, version 23.8.0, ChemAxon, https://www.chemaxon.com), which was then embedded in three dimensions with CORINA 5.0 (Sadowski et al. 1994; Schwab 2010), generating a single starting conformation. Protein and ligand MOL2 files were obtained from PDB structures using the SPORES (ten Brink & Exner 2009) program without altering the assigned protonation states. Molecular docking was performed with PLANTS (Korb et al. 2009) (v1.2) employing the PLANTSCHEMPLP scoring function, a search speed setting of speed 1, and binding site parameters defined as center coordinates (30.540, 45.340, 35.050) with a radius of 12Å. For each receptor, several combinations of flexible active-site side chains were tested, and those reported here are those that gave consistent poses on visual inspection. His 344 was treated as flexible in the At2g17340 model and Lys386 in the Chlamydomonas model. Residues are given in the numbering of each respective structure. The docking output was clustered (maximum 10 structures, cluster RMSD = 2Å), and the resulting complexes were visually inspected for final selection.

### Virtual Screening

Six substructure queries were derived from the parent compound by progressive simplification, from retaining the full parent structure of RMDPi-1 to progressively retain only the 2,2’-Bipyridine core and were used as query for a substructure search over two commercial compound collections: the Enamine In-Stock library (approximately 4 million compounds) and the Enamine REAL database (approximately 5 billion compounds). The substructure queries were organised into nested tiers to stratify the retrieved compounds by their degree of similarity to the parent compound. The rationale was twofold: to identify close analogues of the parent that preserve O,O-chelation, and to sample more broadly around the biaryl scaffold. Every molecule containing the query as a substructure was retained as a superstructure candidate. The union of all hit lists, after deduplication, comprised 2,462 superstructures from the Enamine In-Stock library and 154,532 superstructures from the Enamine REAL database, which together constituted the screening set for the docking campaign against AtRMDP1. The retrieved superstructures were prepared for docking by generating three-dimensional coordinates from the SMILES strings. For every input molecule, the predominant protomer and tautomer at physiological pH were enumerated with the ChemAxon cxcalc utility at pH 7.4. Each protomer was then embedded in three dimensions with CORINA 5.0 (Sadowski et al. 1994; Schwab 2010), generating a single starting conformation. The prepared compounds were docked into the crystallographic coordinates of At2g17340 as described for RMDPi-1. Compounds were first ranked by docking score and then subjected to visual inspection of the predicted binding mode, with priority given to poses that reproduced coordination of the catalytic Mg^2+^ ion. Chemical diversity, synthetic accessibility and physicochemical properties were also considered. On this basis, 14 compounds (Supplementary Files S1 and S2) were prioritised, purchased from Enamine and evaluated experimentally.

### *Chlamydomonas reinhardtii* strains, growth conditions, transformation and cleavage assay

The *srta1* strain (Zou et al. 2026b) was used as the control background in this study. *rmdp* mutants were generated using targeted insertional mutagenesis using the CRISPR/Cas9-based method (Picariello et al. 2020). For complementation, the plasmid pCM2-YZ208 was constructed to contain the native promoter (2,244 bp upstream of the start codon), the genomic DNA sequence of *CrRMDP* fused to the *mVenus* coding sequence, and the PS AD terminator. The Chlamydomonas cells were grown in tris-acetate-phosphate (TAP) medium at 22°C under a 16 h light/8 h dark photoperiod (LD 16:8). Transformations were carried out using glass bead method (Kindle 1990). For Chlamydomonas GFP-cleavage assays, cells at a density of 2*10^6^ cells/ml were used. Cells in 1 ml culture were treated with 50 µM RMDPi-1 for 4h and harvested with two consecutive spins: the first to remove most of the supernatant and the second to compact the cell pellet. Pellets were resuspended in 50 pl of 2*Laemmli sample buffer and heated at 95°C for 5 min. Samples were then vortexed for 30 s and centrifuged at 17,000\**g* for 10 min to pellet cellular debris. The resulting supernatants were used for SDS-PAGE analysis. For oxidative stress growth assays, cells were precultured to the early stationary phase, and equal numbers of cells (2.5 10^5^) were spotted onto TAP agar plates supplemented with 0.4 µM MV. Cell growth was documented after 30 days of incubation.

### Vector construction and plant transformation

All primers and vectors used in this study are listed in Supplementary file S8. To generate knockout mutants for each candidate gene using the CRISPR-Cas9 system, six guide RNAs (gRNAs) were designed. Two gRNAs targeting *AT2G17320* were cloned into the gRNA expression cassettes pGG-A-AtU6-26-BbsI-ccdB-BbsI-B and pGG-B-AtU6-26-BbsI-ccdB-BbsI-C. Two gRNAs targeting *AT2G17340* were introduced into pGG-C-AtU6-26-BbsI-ccdB-BbsI-D and pGG-D-AtU6-26-BbsI-ccdB-BbsI-E and two gRNAs targeting *AT4G35360* were introduced into pGG-E-AtU6-26-BbsI-ccdB-BbsI-F and pGG-F-AtU6-26-BbsI-ccdB-BbsI-G. The six gRNA expression cassettes were assembled into the destination vector pFASTRK-AtCas9-AG using GreenGate cloning. This vector enables selection of transformed seeds based on red fluorescent seed coat expression. The resulting constructs were transformed into *Agrobacterium tumefaciens* strain GV3101 (Larebeke et al. 1974) via freeze-thaw using liquid nitrogen. Wild-type *(WT)* Arabidopsis plants, pHusion marker lines expressing 35S::RFP-GFP-Atg8a and *atg5/7* deficient pHusion marker lines were transformed using the floral dip method as previously described (Clough & Bent 1998). Transgenic T0 seeds displaying red seed coat fluorescence of each genetic background were selected and plated. The one week old seedlings were subjected to a 37°C heat treatment for 24 hours according to Kurokawa et al. (2021) and transferred to soil. T0 plants were genotyped by PCR, and targeted mutations were confirmed by sequencing. For each confirmed T0 line, ten non-fluorescent T1 seeds lacking the Cas9 construct were selected and grown in soil. Homozygous T1 knockout plants were identified by PCR and confirmed by sequencing, after which plants were harvested for T2 seed collection.

### Methyl viologen (MV)-induced cell death assay

Arabidopsis seedlings were grown under standard growth conditions. Six-day-old seedlings were treated in 0.5 MS medium with 2.5 µM RMDPi-1 for 4 h and subsequently transferred to control medium or medium supplemented with 100 nM MV on plates. After 7 days of growth on MV-containing plates, seedlings were imaged using confocal laser scanning microscopy (CLSM). For imaging, seedlings were mounted in standard 0.5*MS medium supplemented with a 1:100 dilution of a 1 mg/ml propidium iodide (PI) stock solution (water-based). Z-stack images were acquired using a Nikon AX-R confocal microscope (galvo scan mode).

### Superoxide staining assay

Six-day-old seedlings were treated in liquid MS medium containing either 0.1% DMSO (control), 100 µM RMDPi-1, or 5 µM MV for 2 h. Following treatment, seedlings were incubated in MS medium supplemented with 30 µM dihydroethidium (DHE; Sigma-Aldrich, D7008) for 15 min in the dark to detect superoxide accumulation. Seedlings were subsequently washed three times with MS medium prior to imaging. DHE fluorescence (excitation/emission: 510/595 nm) was detected using confocal microscopy (Nikon). Fluorescence intensity measurements were quantified using ImageJ.

### Metabolomics sample preparation

#### Arabidopsis

One-week-old seedlings were transferred to 6-well plates containing 3 ml half-strength MS medium and treated for 6 h with either DMSO or 20 µM RMDPi-1. Seedlings were rinsed twice in freshly prepared, filtered 100 mM PBS, pH 7.4. Each replicate comprised 40 seedlings (∼50 mg frozen material).

#### Chlamydomonas

Cultures of the srta (control) and *rmdp* lines were inoculated (0.5:20 ratio) the day before treatment and treated for 24 h in shaker flasks (10 ml each), with an aliquot taken for cell counting. Cells were harvested by centrifugation (500xg, 5 min), rinsed twice in freshly prepared, filtered 100 mM PBS (pH 7.4), resuspended in 1 ml, transferred to screw-cap tubes, and pelleted again (500xg, 3 min) before removal of the supernatant.

#### Metabolite extraction (both species)

Frozen samples were homogenised in a FastPrep-24 instrument (MP Biomedicals; 5 m/s, 30 s, 3 cycles). Pre-chilled (-20°C) 70% methanol (HPLC grade, 1 ml) was added, followed by a further homogenisation step (5 m/s, 30 s) and incubation at -20°C for 1 h. Extracts were cleared by centrifugation (13,300 rpm, 30 min, 2°C). A 20 µl aliquot was reserved for LC-MS analysis; a further 850 µl was dried to completion in a vacuum concentrator and resuspended in 375 µl Milli-Q water plus 150 µl 0.4 M sodium phosphate buffer (pH 7) for NMR analysis.

### NMR metabolomics analysis

^1^H NMR spectra were acquired at 298 K using a Bruker Avance III 600 MHz spectrometer (Bruker BioSpin, Rheinstetten, Germany) equipped with a cryogenic probe and an autosampler. Data were collected using the Bruker zgesgp pulse sequence with 128 scans, an acquisition time of 1.8 s, and a relaxation delay of 4 s. A total of 65,536 data points were acquired over a spectral width of 17,942.584 Hz. All spectral processing was performed using Chenomx NMR Suite (version 7.1; Chenomx Inc., Edmonton, AB, Canada). Spectra were manually phase- and baseline-corrected and referenced to the TSP methyl resonance at 0.00 ppm. The processed spectra were subsequently divided into 0.01-ppm buckets using Chenomx, and the resulting bucketed data were used for univariate and multivariate statistical analyses. Metabolites were annotated using the Chenomx NMR Suite Profiler, the Human Metabolome Database (HMDB), and previously published literature. Where required, metabolite assignments were further supported by two-dimensional NMR experiments, including COSY and TOCSY.

### LC-MS metabolomics analysis

Chromatographic separation in negative ionization mode was performed using an ACQUITY Premier BEH Amide or C18AX column (1.7 pm, 2.1 * 100 mm) maintained at 40°C, with a flow rate of 0.35 ml/min and an injection volume of 5 µl. The aqueous mobile phase (A) consisted of 15 mM ammonium acetate adjusted to pH 9.0, and mobile phase B was acetonitrile.

The gradient started at 95% B, decreased to 85% B at 2 min, 60% B at 10 min, and 50% B at 13 min. The initial condition of 95% B was restored at 13.1 min and maintained until 16 min for column re-equilibration. LC-MS data were processed using XCMS. Chromatographic peaks were detected using the CentWave algorithm with 5 ppm mass tolerance, snthresh = 10, and two peak-width settings (3-10 s and 4-15 s) to accommodate differences in chromatographic peak widths. Retention times were corrected using Obiwarp, peaks were grouped across samples using PeakDensity, missing peaks were filled, and integrated peak areas were exported as the final feature table for statistical analysis. Features corresponding to metabolites of interest, particularly phosphorylated sugars and related compounds, were selected from the XCMS output based on their accurate *m/z* values and subjected to statistical analysis. Metabolite annotation was then performed using authentic standards and, where possible depending on metabolite concentration, MS/MS fragmentation data, considering mass accuracy, retention time, and fragmentation patterns.

### Statistical Analysis and Image Preparation

Statistical analysis was performed using GraphPad Prism (version 10.0.1, GraphPad Software) unless otherwise stated. Figures were prepared using Adobe Photoshop and IllustratorCC (Adobe). Adjustments to brightness and contrast were applied equally to corresponding images within figures.

## Acknowledgments and Funding

Funding for AND was provided by Formas - a Swedish Research Council for Sustainable Development (Grant #2023-00756; 2019-01565); The Carl Tryggers Foundation (#CTS20:93); Epic-XS (#0000215); Chemical Biology Consortium of Sweden (CBCS) Large Project Support (#LP23F:004). This project utilised the NMR infrastructure at the Swedish University of Agricultural Sciences (SLU), funded by the Faculty of Natural Resources and Agricultural Sciences and the Swedish Research Council (grant no. 2021-00167). The Chemical Biology Consortium Sweden (CBCS), node KI, is a national research infrastructure funded by the Swedish Research Council (dr.nr. 2021-00179) and SciLifeLab. This work was supported by the Knut and Alice Wallenberg Foundation (grant 2018.0026 to P.V.B), the Swedish Research Councils VR (grant 2019-04250 to P.V.B. and E.A.M) and Formas (grant 2017-00541 to P.V.B and 2021-01812 to E.A.M), the Carl Tryggers Foundation (grant 23:2470 to I.S.), and the Crops for The Future research programme at the Swedish University of Agricultural Sciences.

The authors declare no conflict of interest.

---

**Supplementary file S1:** RMDPi-1 analog structures and autophagic responses

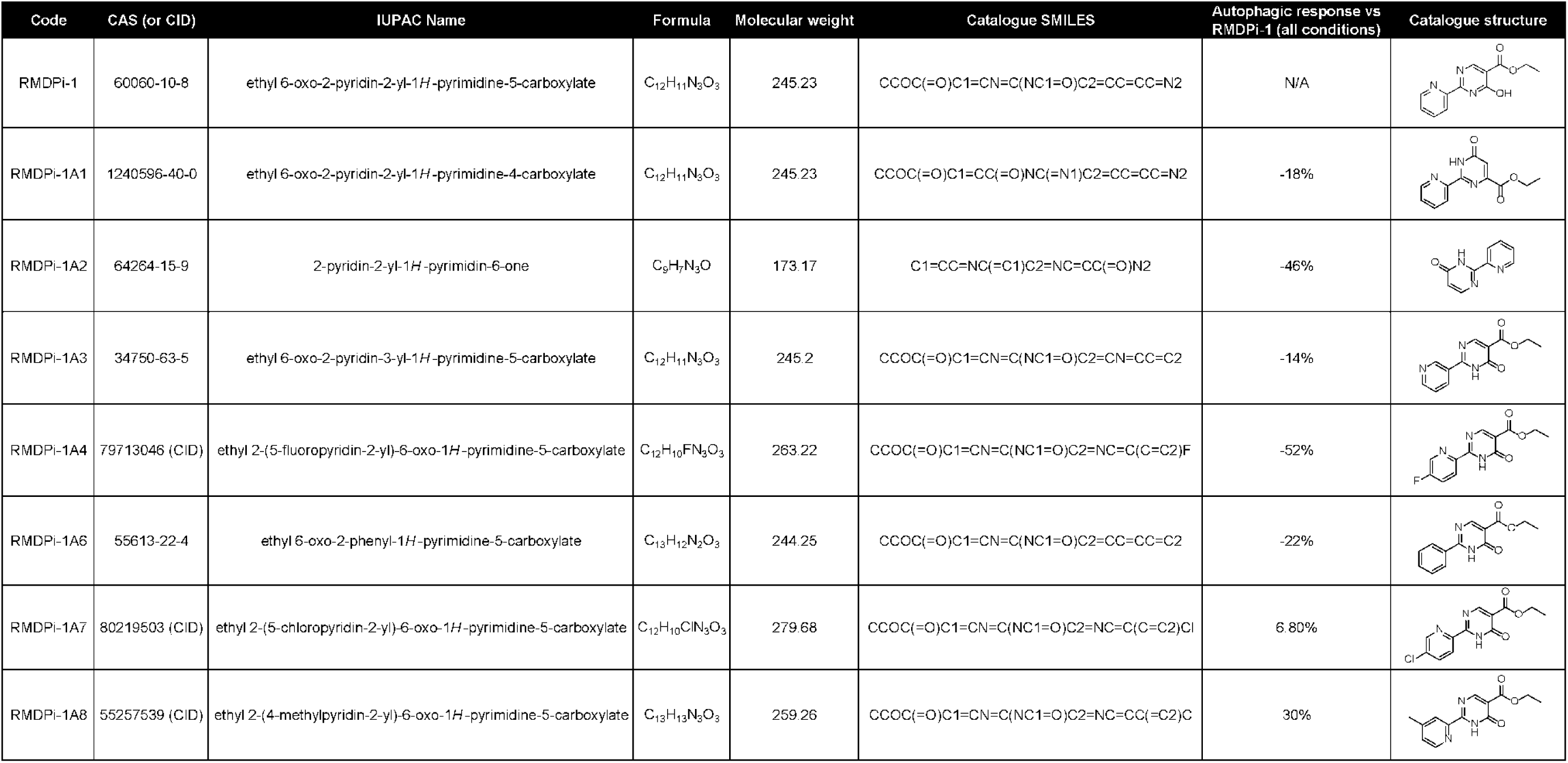

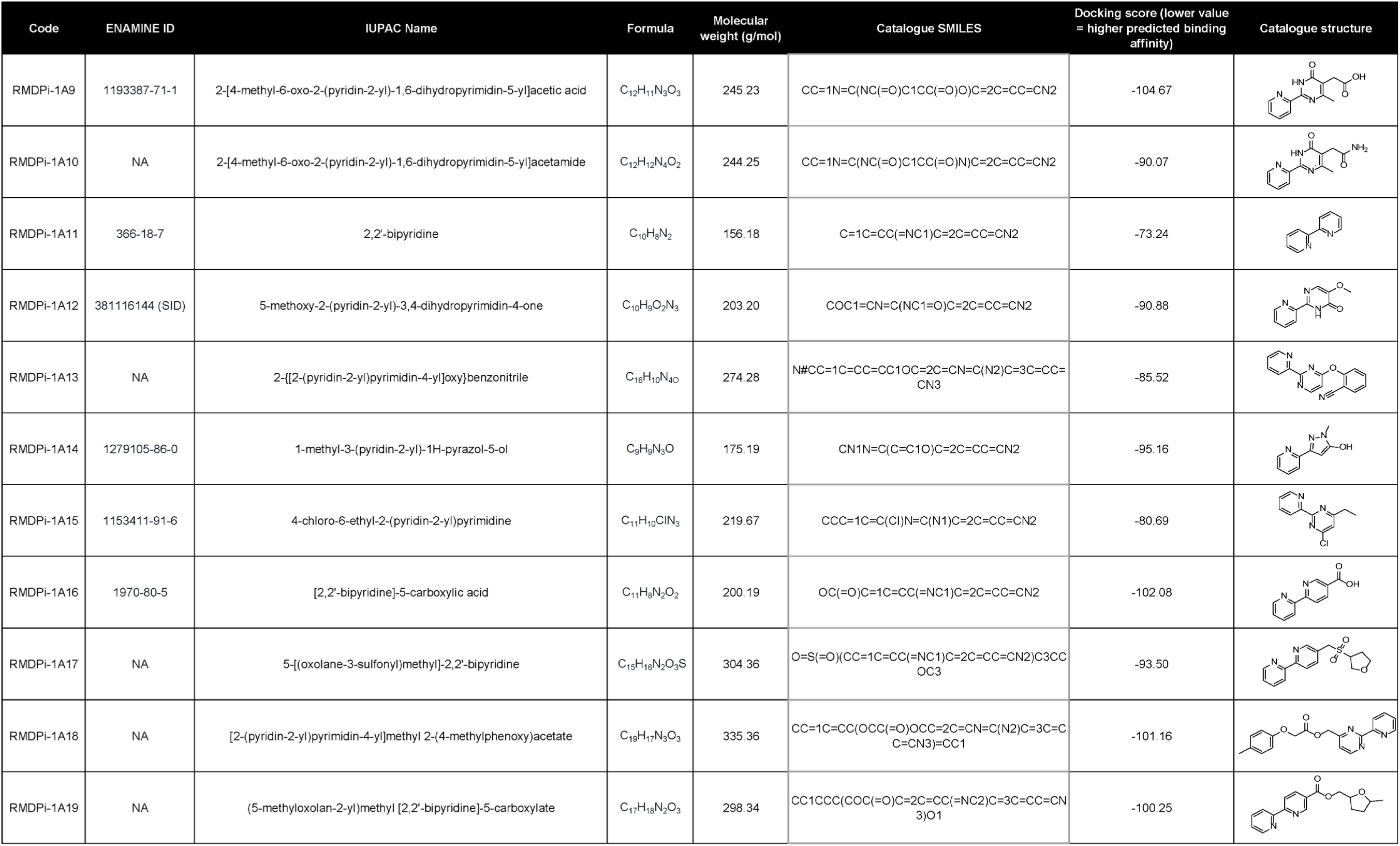

**Supplementary file S2:** Chemical synthesis details

## RMDPi-1A6: Ethyl 6-oxo-2-phenyl-1,6-dihydropyrimidine-5-carboxylate

Diethyl ethoxymethylenemalonate (237.8 mg, 1.1 mmol) and sodium hydroxide (44.0 mg, 1.1 mmol) were added to benzimidamide hydrochloride hydrate (174.6 mg, 1.0 mmol) in 1.5 ml of ethanol. The reaction mixture was refluxed with stirring at 100 °C (sand bath) for 4 hr. LC/MS indicated the reaction was finished. The mixture was cooled to room temperature, and the precipitate was collected by filtration and washed 2-3 times with ethanol (95%) to give 145.0 mg of white needle crystals with 59% yield. ^1^H NMR (CDCl3): 1.42 (t, *J* = 7.1Hz, 3H), 4.44 (q, *J* = 7.1Hz, 2H), 7.54-7.61 (m, 3H), 8.39-8.42 (m, 2H), 8.94 (s, 1H), 13.00 (br., 1H); ^13^C NMR (CDCl3): 14.46, 61.78, 112.36, 128.80, 129.17, 132.45, 133.12, 160.70, 163.32, 165.12; LC-MS: calculated for C13H13N2O3: [M + H]^+^, 245.09; found, 245.2.

## RMDPi-1A7: Ethyl 2-(5-chloropyridin-2-yl)-6-oxo-1,6-dihydropyrimidine-5-carboxylate

Diethyl ethoxymethylenemalonate (76.8 mg, 0.355 mmol) and sodium methoxide (19.2 mg, 0.355 mmol) were added to 5-chloropicolinimidamide (68.2 mg, 0.355 mmol) in 2.0 ml of methanol. The reaction mixture was heated with stirring at 80 °C (sand bath) overnight. The mixture was cooled to room temperature and the precipitate was collected by filtration and purified with HPLC (acetonitrile to water=20-100% /25 min. with 0.005% of formic acid, flowrate=20.0 ml/min., detection wavelength=330nm.) to give 60.0 mg of product in 60% of yield after lyophilization. ^1^H NMR (DMSOd6): 1.28 (t, *J* = 7.1Hz, 3H), 4.25 (q, *J* = 7.1Hz, 2H), 8.21 (dd, *J* = 8.5Hz, *J* = 2.4Hz, 1H), 8.35 (d, *J* = 8.5Hz, 1H), 8.52 (s, 1H), 8.84 (d, *J* = 2.1Hz, 1H), 12.87 (br., 1H); ^13^C NMR (DMSOd6): 14.13, 60.52, 124.56, 134.77, 137.94, 146.70, 148.04, 163.49; LC-MS: calculated for C12H11ClN3O3: [M + H]^+^, 280.05; found, 280.1.

## RMDPi-1A8: Ethyl 2-(4-methylpyridin-2-yl)-6-oxo-1,6-dihydropyrimidine-5-carboxylate

Diethyl ethoxymethylenemalonate (283.2 mg, 1.31 mmol) and sodium methoxide (70.8 mg, 1.31 mmol) were added to 4-methylpicolinimidamide hydrochloride (224.8 mg, 1.31 mmol) in 2.0 ml of methanol. The reaction mixture was refluxed with stirring at 80 °C (sand bath) overnight. LC/MS indicated the reaction was completed. The mixture was cooled to room temperature and the precipitate was collected by filtration and washed 2-3 times with ethanol to give 252.0 mg of crude product. 100.0 mg of the product was purified with HPLC (acetonitrile to water = 20-100%/25 min. with 0.005% formic acid, flowrate = 20.0 ml/min, detection wavelength = 320nm.) to give 76.0 mg of product after lyophilization. ^1^H NMR (CDCl3): 1.40 (t, *J* = 7.1Hz, 3H), 2.50 (s, 3H), 4.41 (q, *J* = 7.1Hz, 2H), 7.37 (d, *J* = 3.4Hz, 1H), 8.33 (s, 1H), 8.56 (s, 1H), 8.81 (s, 1H), 11.26 (br., 1H); ^13^C NMR (CDCl3): 14.41, 21.39, 61.60, 124.18, 128.47, 146.52, 148.93, 157.50, 160.21, 163.90; LC-MS: calculated for C13H14N3O3: [M + H]^+^, 260.10; found, 260.2.

## RMDPi-1 probe: Ethyl 2-(4-((2-(3-(but-3-yn-1-yl)-3*H*-diazirin-3-yl)ethyl)carbamoyl)phenyl)-6-oxo-1,6-dihydropyrimidine-5-carboxylate

A solution of 4-(5-(ethoxycarbonyl)-6-oxo-1,6-dihydropyrimidin-2-yl)benzoic acid (25.7 mg, 0.089 mmol), HBTU (44.0 mg, 0.116 mmol) and DIEA (40 µl, 0.232 mmol) in DMF (10 ml) was stirred for 10 min at room temperature then 2-(3-(but-3-yn-1-yl)-3*H*-diazirin-3-yl)ethan-1-aminium methanesulfonate (25.0 mg, 0.107 mmol) was added. The reaction mixture was stirred at room temperature for 6 hr. The DMF was removed under vacuum (60 °C). Methanol was added to the residue and the mixture was centrifuged to remove precipitates. The methanol solution was purified by preparative HPLC (acetonitrile to water =20-80%/25min. with 0.005% formic acid, flow rate = 20.0 ml/min., detection wavelength = 230 nm.) to give 6.3 mg of product at 17% yield after lyophilization. ^1^H NMR (DMSOd6): 1.29 (t, *J* = 7.1Hz, 3H), 1.64 (t, *J* = 7.4Hz, 2H), 1.68 (t, *J* = 7.1Hz, 2H), 2.02 (dt, *J*= 2.6Hz, *J* =7.4Hz, 2H), 2.83 (t, *J* = 2.6Hz, 1H), 3.17-3.20 (m, 2H), 4.27 (q, *J* = 7.1Hz, 2H), 7.78 (d, *J* =8.5Hz, 2H), 8.25 (d, *J* =8.1Hz, 2H), 8.66 (s, 1H), 8.66 (t, *J* = 5.4Hz, NH), 13.28 (s, 1H); ^13^C NMR (DMSOd6): 12.70, 14.16, 27.30, 31.27, 31.82, 34.40, 34.52, 60.52, 71.82, 83.15, 127.45, 128.42, 137.64, 137.69, 158.95, 163.48, 165.22, 165.30, LC-MS: calculated for C21H22N5O4: [M + H]^+^, 408.17; found, 408.3.

## Control Probe: *N*-(2-(3-(But-3-ynyl)-3*H*-diazirin-3-yl)ethyl)benzamide

Benzoic acid (10.0 mg, 0.082 mmol), 1-Hydroxybenzotriazole (HOBt) (16.6 mg, 0.123 mmol), and 1-ethyl-3-(3-dimethylaminopropyl)carbodiimide (EDCI) (19.1 mg, 0.123 mmol) were dissolved in 0.5 ml of dry DMF, then triethylamine (22.8 µl, 0.164 mmol) was added. The mixture was stirred for 10 min. 3-(3-butyn-1-yl)-3*H*-diazirine-3-ethanamine (12.4 mg, 0.090 mmol) was added to the reaction mixture and stirred overnight at room temperature. LC/MS indicated that the starting material was consumed. 0.3 ml of methanol was added to the reaction mixture and purified with semi-preparative HPLC (acetonitrile to water = 10-80%/25min. with 0.005% formic acid, flow rate = 7.0 ml/min., detection wavelength = 230 nm.) to give 16.5 mg of white solid at 84% yield after lyophilization. ^1^H NMR (CDCl3): 1.68 (t, *J* = 7.3Hz, 2H), 1.82 (t, *J* = 6.6Hz, 2H), 1.98 (t, *J* = 2.6Hz, 1H), 2.03 (dt, *J* = 2.6Hz, *J* = 7.2Hz, 2H), 3.31 (dt, *J* = 6.3Hz, *J* = 6.5Hz, 2H), 6.34 (s, 1H), 7.42-7.45 (m, 2H), 7.49-7.51 (m, 1H), 7.78-7.79 (m,2H). ^13^C NMR (CDCl3): 13.32, 27.06, 32.25, 32.62, 35.02, 69.57, 82.81, 127.02, 128.75, 131.71, 134.52, 167.67. LC-MS: calculated for C14H16N3O: [M + H]^+^, 242.13; found, 242.2.

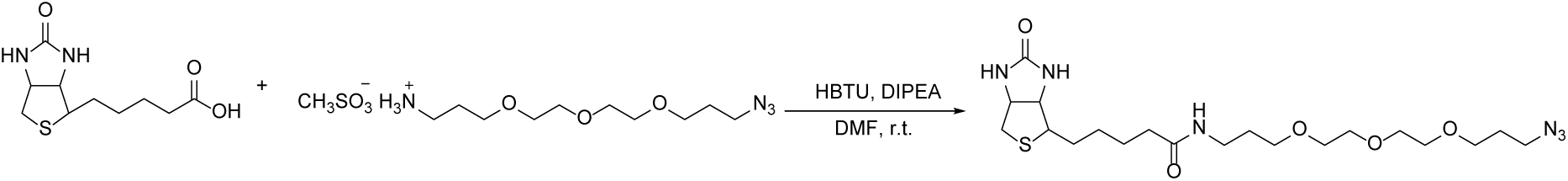

## Biotin tag: *N*-(3-(2-(2-(3-Azidopropoxy)ethoxy)ethoxy)propyl)-5-(2-oxohexahydro-1*H*-thieno[3,4-*d*]imidazol-4-yl)pentanamide

A solution of biotin (189.2 mg, 0.775 mmol), HBTU (318.2 mg, 0.839 mmol) and DIEA (290 µl, 1.678 mmol) in DMF (10 ml) was stirred for 10 min at room temperature before adding to a solution of 3-(2-(2-(3-azidopropoxy)ethoxy)ethoxy)propan-1-aminium triacetate in DMF (6.0 ml). The reaction was stirred overnight at room temperature, after which the DMF was removed by vacuum, resulting in an oil. The residue was purified with HPLC (acetonitrile to water = 20-100%/25min. with 0.005% formic acid, flow rate = 20.0 ml/min., detection wavelength = 220 nm.) to give 232.0 mg of product at 76% yield after lyophilization. ^1^H NMR (CD3OD): 1.42-1.48 (m, 2H), 1.58-1.78 (m, 6H), 1.80-1.86 (m, 2H), 2.20 (t, *J* = 7.4Hz, 2H), 2.71 (d, *J* = 12.7Hz, 1H), 2.93 (dd, *J* = 5.0Hz, *J* = 12.7Hz, 2H), 3.19-3.23 (m, 1H), 3.26 (t, *J* = 6.8Hz, 2H), 3.40 (t, *J* = 6.7Hz, 2H), 3.52 (t, *J* = 6.1Hz, 2H), 3.56 (t, *J* = 6.2Hz, 2H), 3.59-3.61 (m, 4H), 3.63-3.65 (m, 4H), 4.30 (dd, *J* = 4.5Hz, *J* = 7.9Hz, 1H), 4.49 (dd, *J* = 4.9Hz, *J* = 7.8Hz, 1H); ^13^C NMR (CD3OD): 26.89, 29.51, 29.81, 30.18, 30.42, 36.85, 37.84, 41.04, 57.01, 61.64, 63.40, 68.92, 69.95, 71.26, 71.30, 71.55, 71.57, 166.11, 175.96; LC-MS: calculated for C20H37N6O5S: [M + H]^+^, 473.25; found, 473.3.

**Supplementary file S3**: Proteomic analysis, contact corresponding author

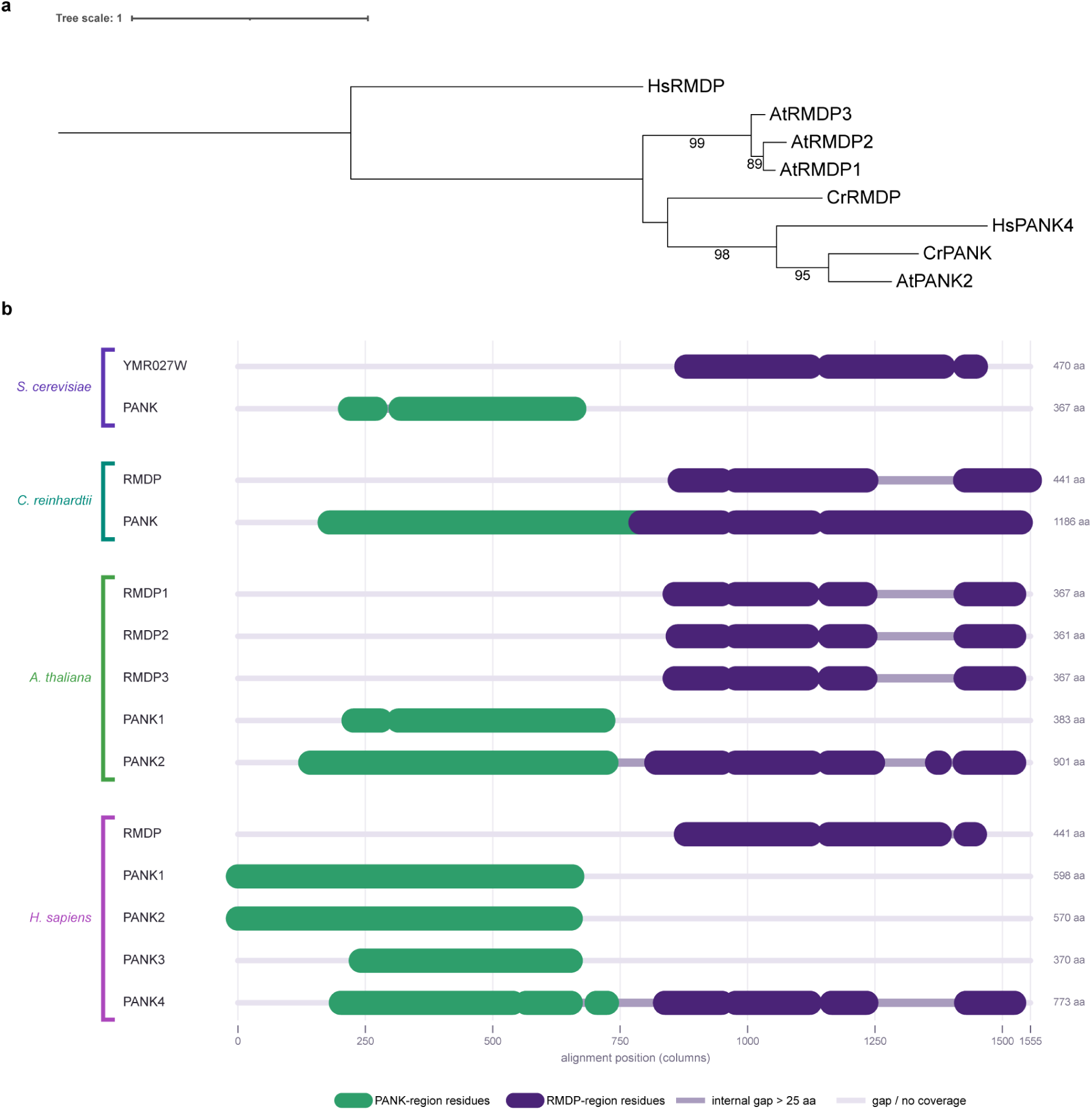

**Supplementary file S4. An overview of subfamily II Domain of Unknown Function 89 (DUF89) / Reactive Metabolite Damage Phosphatase (RMDP) proteins, a,** Maximum-likelihood phylogenetic tree. The sequences were aligned using MAFFT L-INS-i v7.490 (Katoh and Standley 2013), and positions with more than 50% gaps were removed using trimAL v1.4. rev15 (Capella-Gutierrez et al. 2009). Phylogenies were computed using IQ-TREE v2.1,4-beta (Minh et al. 2020). Phylogenetic models were automatically selected using ModelFinder (Kalyaanamoorthy et al. 2017). Branch support was assessed using ultrafast bootstrap approximation with 1,000 bootstrap replicates (Hoang et al. 2018). **b,** Alignment of RMDPs, PANK and PANK-RMDP domain sequences in several representative species including *Saccharomyces cerevisiae, Chlamydomonas reinhardtii, Arabidopsis thaliana* and *Homo sapiens*

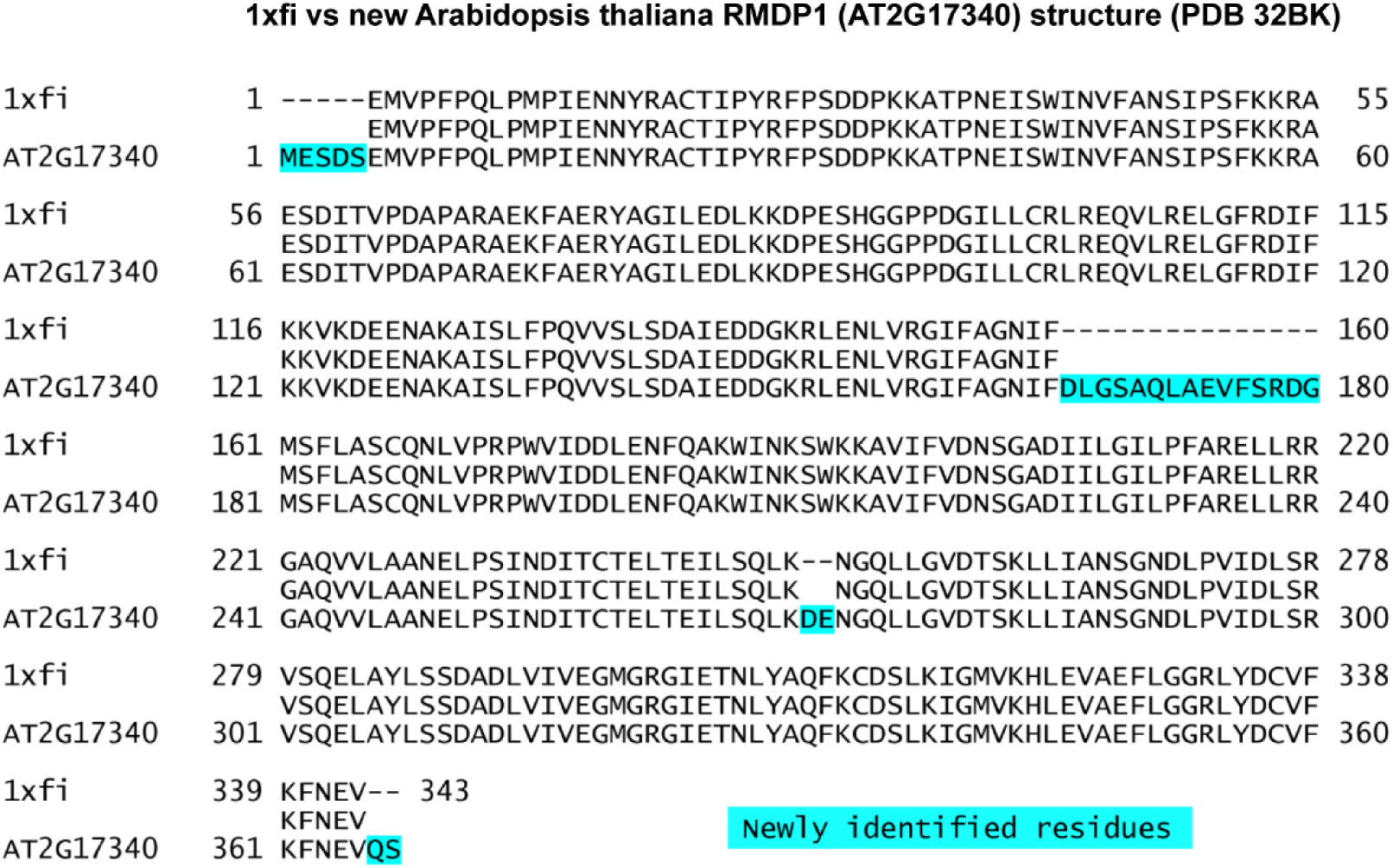

**Supplementary file S5.** Comparison of AtRMDPI structure co-incubated with RMDPi-1A16 with the previously published 1xfi (PDB) structure. Newly releaved residues are marked in cyan

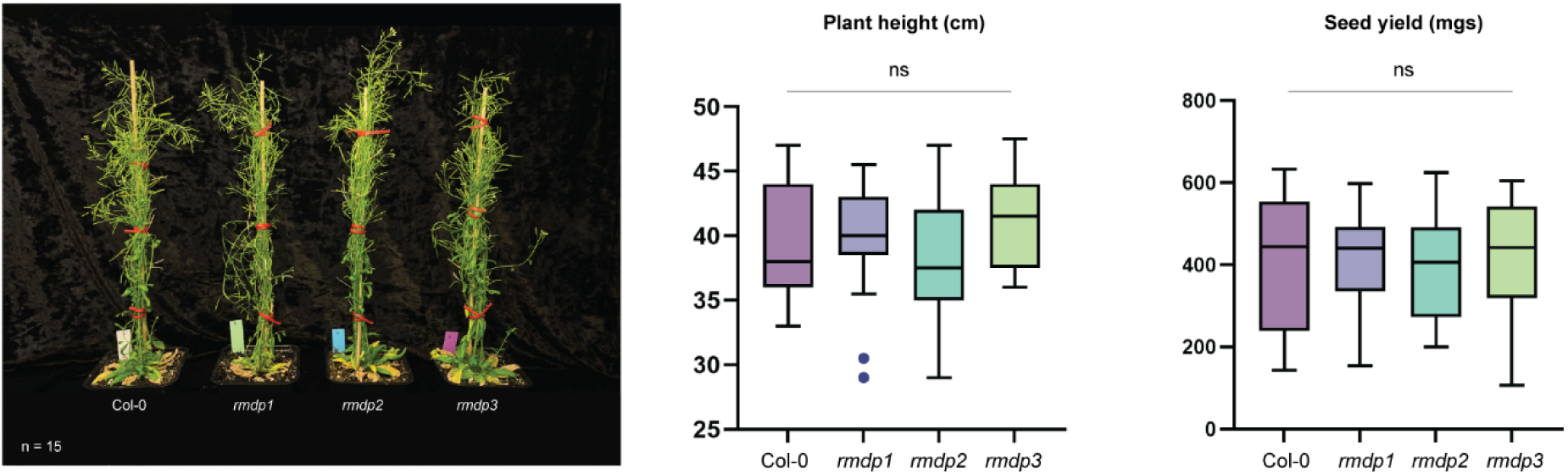

**Supplementary file S6.** Phenotyping of single knockout (T-DNA) lines of the Reactive Metabolite Damage Phospatases (RMDPs) Arabidopsis encodes three RMDP paralogs: At2g17340 (RMDP1), At2g17320 (RMDP2), At4g35360 (RMDP3). One-way ANOVA, ns=non-significant n=15 biological replicates.

**Supplementary file S7**: Metabolome analysis, contact corresponding author

Purity estimation based on integrated signals in the 1H NMR spectrum: >99.3%

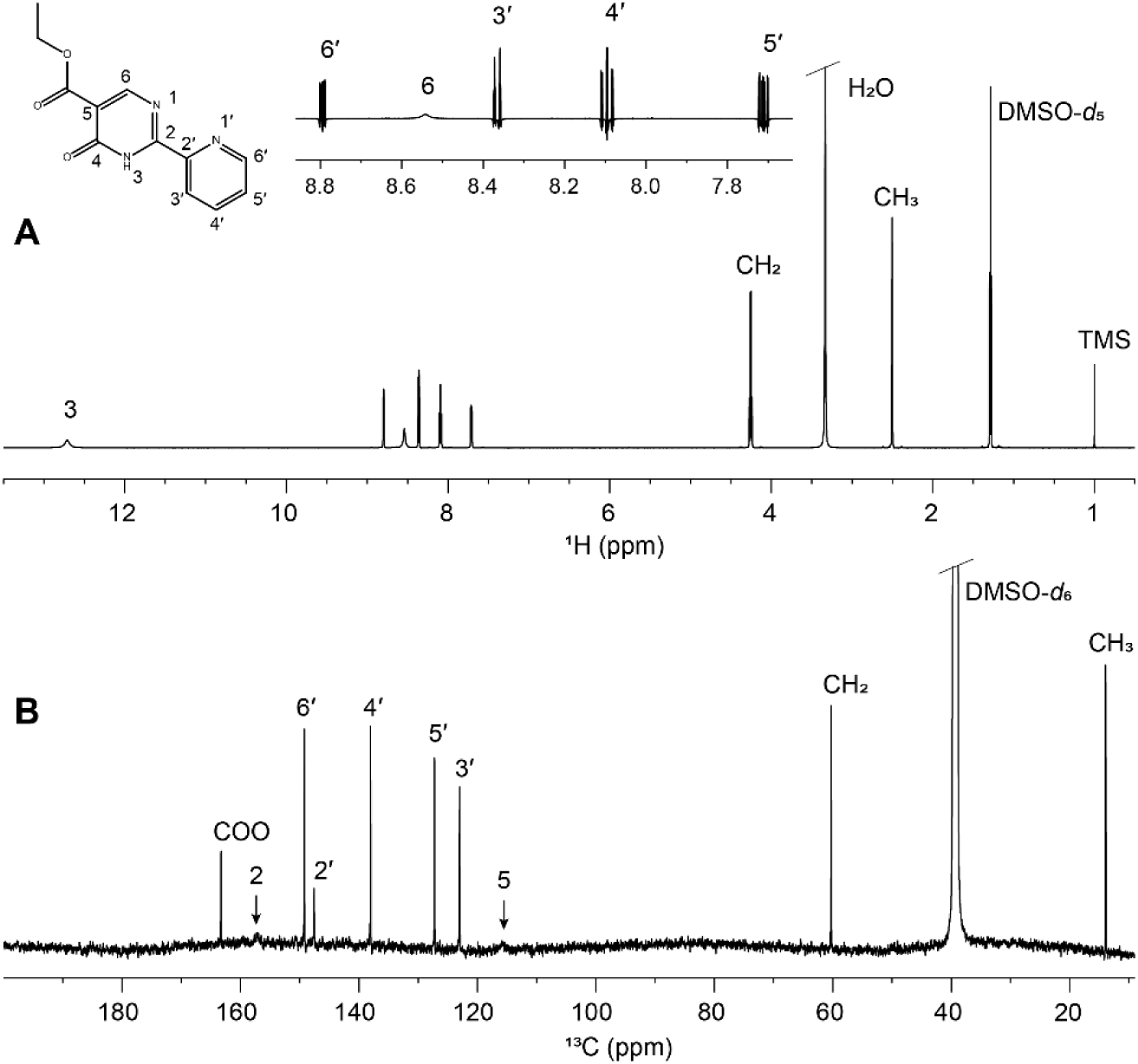

**Supplementary file S8.** A) ^1^H and B) ^13^C NMR spectra of RMDPI-1 with assigned resonances. The numbering is shown on the structural formula of RMDPI-1. The inset shows an expansion of the aromatic region with gaussian processing, which was used for extraction of coupling constants. The signals from H6, C5, and C2 were considerably broadened whereas C4 and C6 were missing. This is probably due to fast exchange between tautomer forms.

**Supplementary file S9:** Full-length coding regions of AtRMDPI (AT2G17340.1)

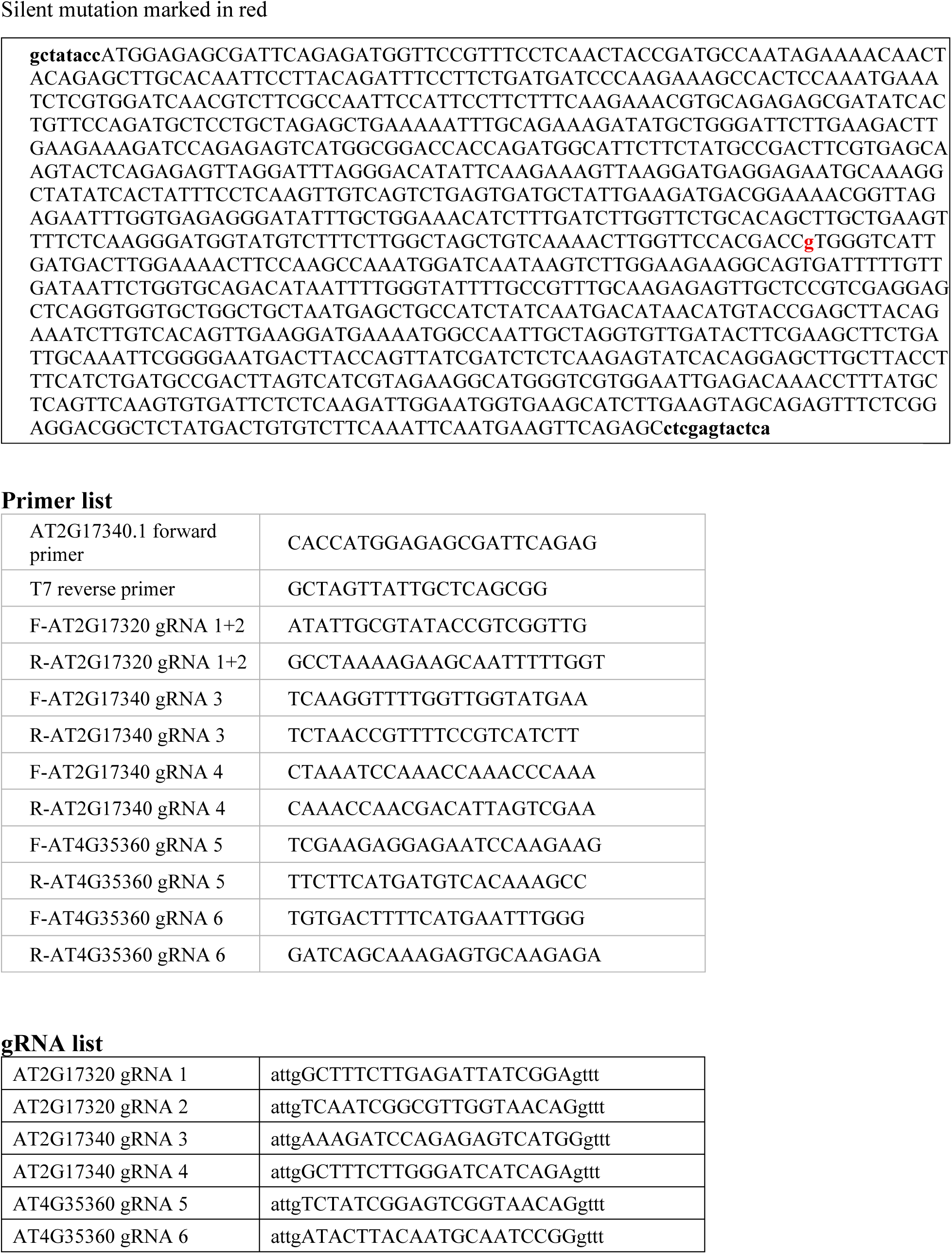

